# A stochastic epigenetic switch controls the dynamics of T-cell lineage commitment

**DOI:** 10.1101/318675

**Authors:** Kenneth K.H. Ng, Mary A. Yui, Arnav Mehta, Sharmayne Siu, Blythe Irwin, Shirley Pease, Satoshi Hirose, Michael B. Elowitz, Ellen V. Rothenberg, Hao Yuan Kueh

## Abstract

Cell fate decisions occur through the switch-like, irreversible activation of fate-specifying genes. These activation events are often assumed to be tightly-coupled to changes in upstream transcription factors, but could also be constrained by *cis*-epigenetic mechanisms at individual gene loci. Here, we studied the activation of *Bcl11b*, which controls T-cell fate commitment. To disentangle *cis* and *trans* effects, we generated mice where two *Bcl11b* copies are tagged with distinguishable fluorescent proteins. Quantitative live microscopy of progenitors from these mice revealed that *Bcl11b* turned on after a stochastic delay averaging multiple days, which varied not only between cells but also between *Bcl11b* alleles within the same cell. Genetic perturbations, together with mathematical modeling, showed that a distal enhancer controls the rate of epigenetic activation, while a parallel Notch-dependent *trans*-acting step stimulates expression from activated loci. These results show that developmental fate transitions can be controlled by stochastic *cis*-acting events on individual loci.

## Introduction

During development, individual cells establish and maintain stable gene expression programs through the irreversible activation of lineage-specifying regulatory genes. A fundamental goal of developmental biology is to understand how and when these activation events are initiated to drive cell fate transitions. The concentrations of active transcription factors in the nucleus are crucial for embryonic patterning and progressive gene expression changes in development (Briscoe and Small, 2015; Davidson, 2010; Jaeger, 2011), and are often assumed to directly dictate rates of target gene transcription (Coulon et al., 2013; Estrada et al., 2016; Phillips, 2015). At the same time, an additional layer of epigenetic control mechanisms acts directly at gene loci on chromosomes, through chemical modification of DNA or DNA-associated histone proteins (Bird, 2002; Tessarz and Kouzarides, 2014), or regulation of chromosome conformation or packing in the nucleus (Felsenfeld and Dekker, 2012). Chromatin modification and accessibility changes are ultimately initiated by the binding and action of *trans*-acting factors; however, while these changes are often assumed to closely follow transcription factor changes, other recent work shows that epigenetic processes could occur slowly (Kaikkonen et al., 2013; Mayran et al., 2018), and could introduce slow, stochastic, rate-limiting steps to gene activation, even when transcription factor inputs are fully present (Berry et al., 2017; Bintu et al., 2016). Despite much work, it has generally remained unclear what role, if any, epigenetic mechanisms play in controlling the timing and outcome of developmental gene activation and cell fate decisions.

Epigenetic control is ordinarily difficult to disentangle from control due to changes in transcription factor activity. However, the two mechanisms can be distinguished by their effects on different gene copies in the same cell (Bonasio et al., 2010). Control due to transcription factor changes occurs in *trans*, and thus affects two copies of the gene in the same cell coordinately; in contrast, epigenetic mechanisms function at single gene copies, in *cis*, and thus could generate distinct activation states for different gene copies in the same cell, a concept that underlies the utility of X-chromosome inactivation and other systems as models for epigenetic gene control (Berry et al., 2015; Deng et al., 2014; Farago et al., 2012; Gendrel and Heard, 2014; Ku et al., 2015; Xu et al., 2006). For this reason, tracking both copies of a gene in the same cell with distinguishable fluorescent proteins can provide insight into the dynamics of *cis* and *trans* regulatory processes (Berry et al., 2015; Elowitz et al., 2002).

Using this approach of tracking two gene copies, we have studied the developmental activation of *Bcl11b*, a key driver of T-cell commitment and identity. To become a T-cell, hematopoietic progenitors transition through a series of developmental states, where they lose alternate lineage potential and eventually commit to the T-cell lineage (Fig. 1A). T-cell lineage commitment requires the irreversible switch-like activation of *Bcl11b*, which serves to repress alternate lineage potential and establish T-lineage identity (Li et al., 2010; Liu et al., 2010). *Bcl11b* is regulated by an ensemble of transcription factors, including Runx1, GATA-3, TCF-1, and Notch, which bind to multiple locations on the gene locus (Li et al., 2013; Kueh et al., 2016). However, even when these developmentally controlled transcription factors have been fully mobilized, *Bcl11b* activation occurs only after an extended time delay of ~4 days, allowing pre-commitment expansion of progenitors (Kueh et al., 2016). During activation, the *Bcl11b* locus remodels its epigenetic state, undergoing changes in DNA methylation and histone modification (Ji et al., 2010; Zhang et al., 2012), nuclear positioning, genome compartmentalization and looping interactions (Hu et al., 2018), and expression of a *cis*-acting lncRNA transcript (Isoda et al., 2017). These observations suggest that the dynamics of *Bcl11b* activation could be determined by epigenetic processes as well as transcription factors.

**Figure 1:**
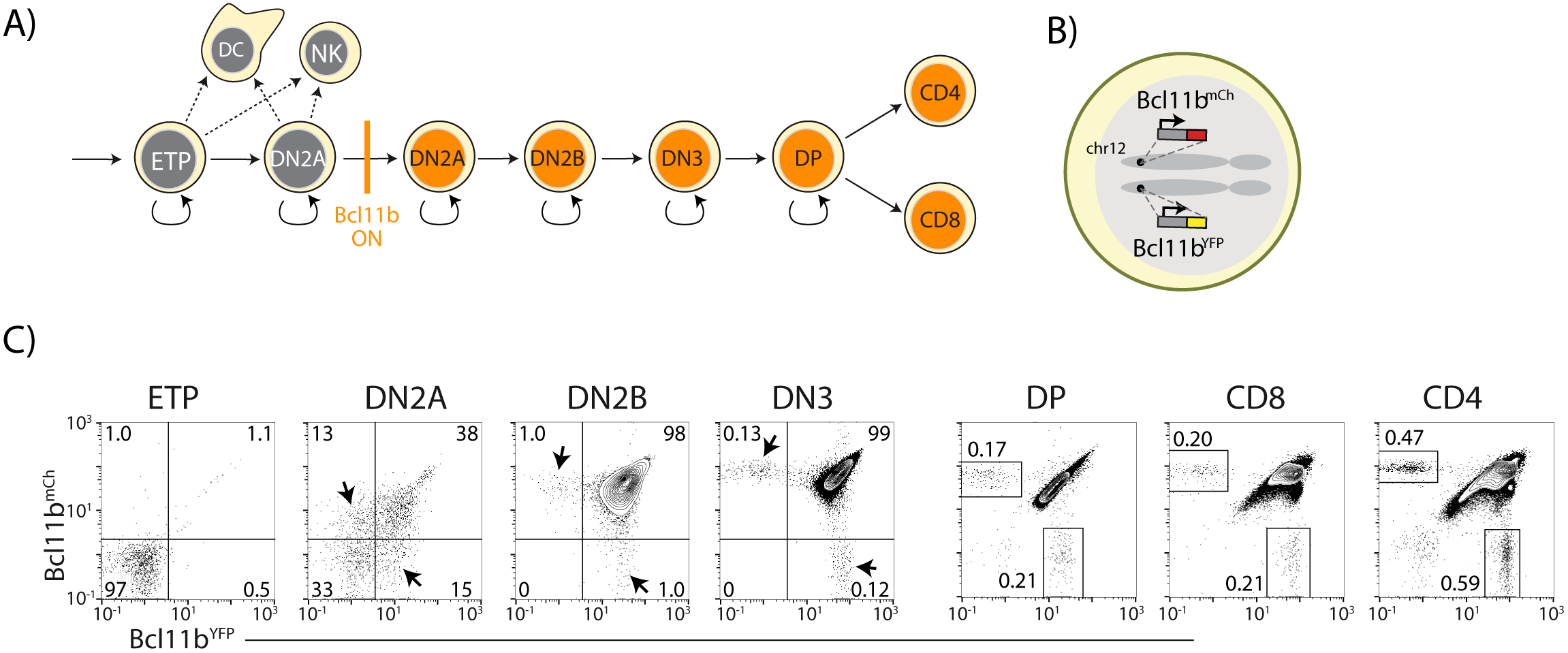
Dual-color *Bcl11b* reporter strategy can reveal epigenetic mechanisms controlling T-cell lineage commitment. A) Overview of early T-cell development. *Bcl11b* turns on to silence alternate fate potentials and drive T-cell fate commitment. ETP – early thymic progenitor; DN2 – CD4^-^ CD8^-^ double negative-2A progenitor; DP – CD4^+^ CD8^+^; NK – natural killer; DC – dendritic cell. B) Dual color *Bcl11b* reporter cells, where two distinguishable fluorescent proteins (YFP and mCherry) are inserted non-disruptively into the same sites on the two chromosomal *Bcl11b* loci. C) Flow cytometry plots show Bcl11b-YFP versus Bcl11b-mCh expression levels in developing T-cell progenitors from dual *Bcl11b* reporter mice. Arrowheads or boxes indicate cells expressing one copy of *Bcl11b*. Results are representative of 3 independent experiments. See also Figure S1.

To separately follow two *Bcl11b* copies in developing cells, we engineered a dual-color reporter mouse, where the two *Bcl11b* copies are tagged with distinguishable fluorescent proteins. We then used quantitative live-cell imaging to follow *Bcl11b* activation dynamics in single progenitor lineages, along with mathematical modeling and perturbation experiments to dissect the relative contributions of *cis*- and *trans*- acting inputs to *Bcl11b* regulation. Our results revealed that activation of *Bcl11b* and consequent T-cell commitment require a stochastic, *cis-*acting epigenetic step on the *Bcl11b* locus. This step occurs independently at the two alleles in the same cell, with a slow timescale spanning multiple days and cell cycles. A separate *trans*-acting step, controlled by the T-cell developmental signal Notch, occurs in parallel with this *cis*-acting step and provides an additional necessary input for *Bcl11b* activation. Finally, we found that over the course of development, T-cell progenitors lose the ability to activate the *cis*-epigenetic switch, and as result, can progress to final differentiated states with only one *Bcl11b* locus stably activated. Together, these results show that intrinsically stochastic events occurring at single gene copies can determine the timing and outcome of mammalian cell fate decisions.

## Results

### Two Bcl11b copies show slow, independent activation in single progenitor lineages

We generated a double knock-in reporter mouse strain, with an mCitrine yellow fluorescent protein (YFP) inserted non-disruptively in the 3’-untranslated region of one *Bcl11b* copy and an mCherry red fluorescent protein (mCh) at the same site in the other copy (Figures 1B and S1). Both *Bcl11b* copies contain a floxed neomycin resistance cassette downstream of the fluorescent protein (Figure S1); however, we have shown conclusively, using Cre-mediated excision, that this cassette has no effect on *Bcl11b* activation (Kueh et al., 2016). We isolated T-cell populations at different stages of development and differentiation directly from dual reporter *Bcl11b* mice, and measured the fraction of cells expressing *Bcl11b* from each allele at stages spanning the initial onset of *Bcl11b* expression (Figure 1C). As reported previously (Kueh et al., 2016; Tydell et al., 2007), *Bcl11b* was inactive in early T-cell progenitors (ETPs), and began to turn on in the subsequent CD4, CD8 double negative (DN)2A stage, becoming expressed in all cells throughout the rest of T-cell development (Figures 1A and 1C). By DN2B and DN3 stages, the large majority of cells had turned on both *Bcl11b* copies. These transitions involve multiple cell cycles each, with about two days between late ETP and DN2A and about three days between DN2A and DN2B. However, in the DN2A compartment where *Bcl11b* gene activation begins, a significant fraction of the cells expressed only one copy of *Bcl11b*, with roughly equal fractions of cells expressing either the YFP or mCherry alleles (Figure 1C, arrowheads). This suggested that the two *Bcl11b* copies could turn on at different times in the same cell during development.

To determine directly whether two *Bcl11b* copies switch on independently in the same cell, we used multi-day timelapse imaging to follow the two *Bcl11b* fluorescent reporters in clonal lineages of developing progenitors. We sorted individual *Bcl11b*-negative DN2A T-cell progenitors from dual *Bcl11b* reporter mice, and cultured them *in vitro* with OP9-DL1 stromal cells (Schmitt and Zúñiga-Pflücker, 2002), confining them in microwells to allow tracking of descendants of each progenitor over multiple days (Figure 2A). The ~1 hour interval between successive frames did not permit complete lineage tracking due to rapid cell movement (Movie S1), but still enabled mapping and visualization of all descendants, and determination of coarse-grained lineage relationships (Figure 2B, C, bottom left). We used OP9-DL1 cells to present the Notch ligand DL1, a critical T-cell developmental signal, and included the supportive cytokines Interleukin (IL)-7 and Flt3 ligand (see Materials and Methods).

**Figure 2:**
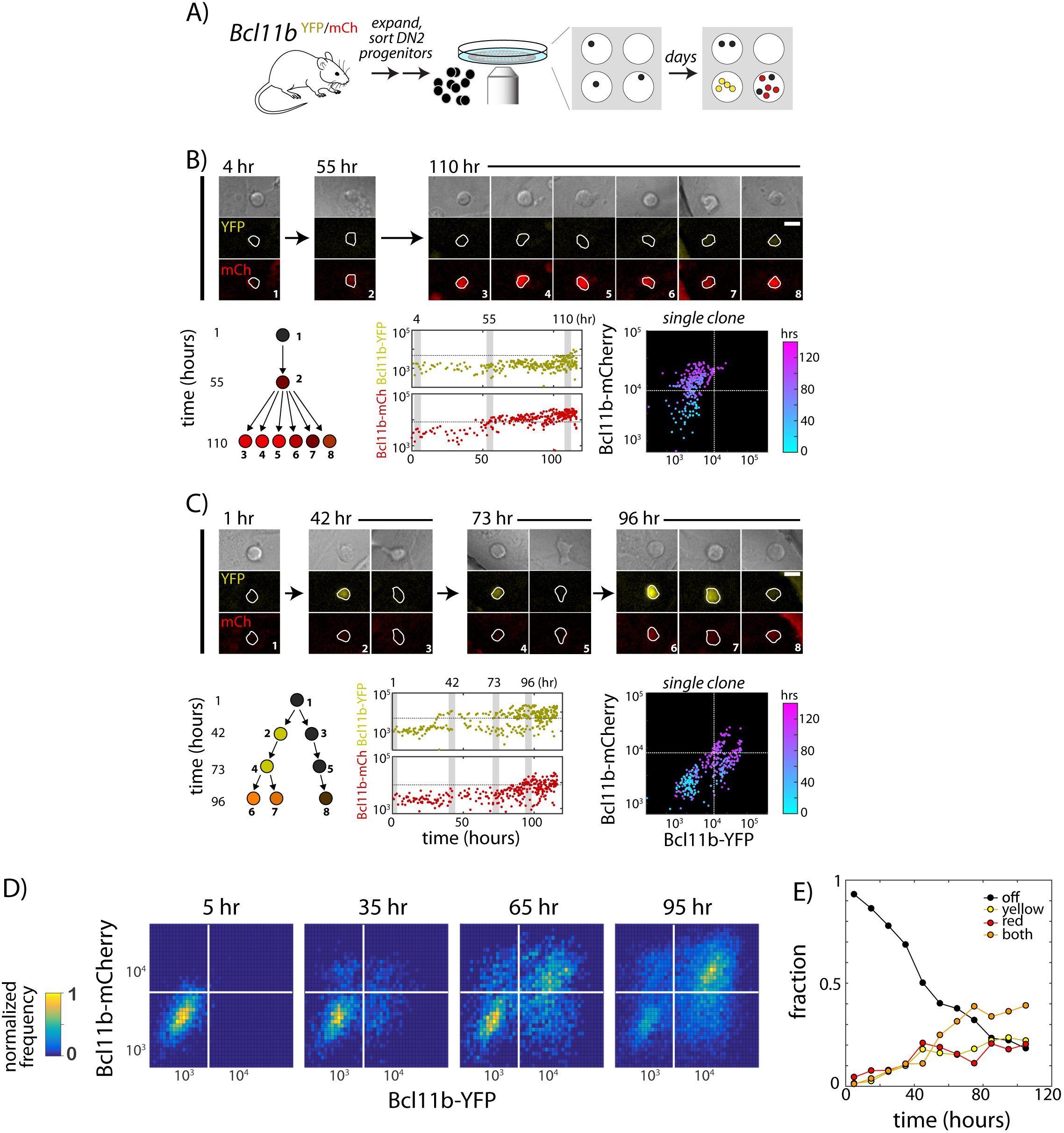
Two copies of *Bcl11b* switch on independently and stochastically in the same cell in single lineages of T-cell progenitors. A-B) *Bcl11b*-negative DN2 cells derived from bone-marrow progenitors were isolated by flow cytometry, cultured within microwells, and followed for 5 days using fluorescence imaging. Cells were then segmented using automated image analysis. B-C). Dynamics of *Bcl11b* activation in two representative clonal progenitor lineages. Timelapse images (top) show developing T-cell progenitors from two representative clones (left), with segmented cell boundaries in white. Numbers (top left) indicate time in hours. Scale bar=10 microns. Trees (bottom left) show coarse-grained cell lineage relationships cells shown here. Plots (center, lower rows) show Bcl11b-YFP and Bcl11b-mCh expression time traces in all cells from a single clone, with vertical gray bars indicating the time points of the image shown on the left. Horizontal lines indicate activation threshold. Colored scatterplots (bottom right) show time evolution of Bcl11b-mCh versus Bcl11b-YFP levels in single clones, from 0h (cyan) to 120 h (purple). D) Heat maps show Bcl11b-YFP and Bcl11b-mCh distributions in the polyconal population at the indicated time points. White lines represent *Bcl11b* expression thresholds. Color bar (left) represents normalized cell numbers at each time point. E) Fractions of cells having different *Bcl11b* allelic expression states, obtained by mixed Gaussian fitting of the heat maps shown. Data represent a cohort of ~200 starting cells from a single timelapse movie. Overall, data show that *Bcl11b* switches on slowly and stochastically in single lineages of progenitors, maintaining alternate activity states in the same clone, heritable across many divisions. Results are representative of 3 independent experiments. See also Figure S2 and Movie S1.

We had previously shown that about three days are required for half of the cells in such DN2A populations to turn on any Bcl11b expression (Kueh et al., 2016). In theory, this delay could reflect requirement for activation of some additional transcription factor. However, even a novel transcription factor would be able to work on both alleles in parallel. Instead, strikingly, imaging revealed strongly asynchronous activation of the two *Bcl11b* copies in the same cell during this time period. Within single clonal lineages of progenitors, one copy of *Bcl11b* could switch on multiple days and cell generations before the other (Figure 2B-C, Movie S1), giving rise to distinct allelic expression states that persisted over multiple divisions. Similar percentages of cells activated Bcl11b-YFP first as compared to those turning on Bcl11b-mCherry first, consistent with independent activation (Figures 2D-E). The results from the same mice also ruled out any imprinting-type allelic bias. Furthermore, in some clones, the times at which a *Bcl11b* allele first turned on differed between progeny of a single cell (Figure 2C, 42 hrs), such that individual progenitors frequently gave rise to clonal descendants with multiple distinct states of *Bcl11b* allelic activation (46.7% heterogeneous after 4d, *N*=15, Figure S2A and S2B). Thus, the allelic bias developed at a clonal level. A substantial percentage (~40%) of all cells remained monoallelic in expression after 4d (Figures 2E), indicating that stochastic locus activation occurs with a slow time constant spanning multiple days. Furthermore, the fractions of cells monoallelically expressing Bcl11b-YFP or Bcl11b-mCh increased with the same dynamics, indicating that each locus is triggered with the same stochastic activation rate. Taken together, these results suggest that timing of the *Bcl11b* activation switch – and the ensuing commitment to become a T-cell – is controlled independently at each *Bcl11b* allele by a stochastic and remarkably slow rate-limiting step.

### A distal enhancer modulates stochastic Bcl11b locus activation

The stochastic transition of *Bcl11b* from an inactive to active state may be controlled by specific *cis*-regulatory DNA elements on the *Bcl11b* locus. Consistent with this idea, we found that graded changes in Notch signaling, GATA-3 activity, and TCF-1 activity alter the likelihood of all-or-none activation, rather than the amplitude of transcription (Kueh *et al.*, 2016). Indeed, in a number of systems, *cis*-regulatory elements do not appear to control transcriptional amplitudes, but instead modulate the probabilities of all-or-none activation (Khan et al., 2011; Walters et al., 1995; Weintraub, 1988). To test how stochastic activation of individual *Bcl11b* alleles may be controlled, we examined the effect of disrupting the one known positive *cis*-regulatory element region, which resides ~850 kb downstream of *Bcl11b* within a “super enhancer” at the opposite end of the same topologically associated domain (Li et al., 2013) (Figure 3A). This region shows distinctive histone marking and some T-lineage-specific transcription factor occupancy even before *Bcl11b* activation (Kueh et al., 2016), lies about 11 kb from the promoter of a *Bcl11b*- associated lncRNA, and loops to the *Bcl11b* gene body in a T-cell lineage specific manner (Hu et al., 2018; Isoda et al., 2017; Li et al., 2013). Like the *Bcl11b* locus itself, this enhancer region is marked by H3K27me3 in non-T lineage cells (Li et al., 2013).

Using standard gene targeting, we deleted this distal ~2kb enhancer region on the Bcl11b-YFP allele, leaving the *Bcl11b-mCherry* allele intact, to generate *Bcl11b*^YFPΔEnh/mCh^ dual reporter mice (Figures 3A and S1), and then analyzed resultant effects on YFP regulation in different T-cell subsets (Figures 3B-C, and S3-S4). These were analyzed either from established young adult *Bcl11b*^YFPΔEnh/mCh^ mice (Figures 3B-C, S3, and S4A-B) or from adult chimeras populated with fetal liver cells from the F_0_ generation (Figure S4). The non-disrupted *Bcl11b-mCherry* allele served as an internal, same-cell control. At the ETP stage, essentially all *Bcl11b* alleles were silent, regardless of whether they had an intact or disrupted enhancer, as expected (Fig. 3B). During the DN2A and DN2B stages, the enhancer-disrupted Bcl11b-YFP allele showed dramatically reduced activation compared to the Bcl11b-mCherry allele in the same cell. Interestingly, at later developmental stages in the thymus and in peripheral T-cell subsets (CD4, Treg, CD8) a large fraction of cells showed expression of the enhancer-disrupted YFP allele, along with the wild type Bcl11b-mCherry allele (Figures 3B-C, S3-4), indicating that the targeted element is not indispensable for *Bcl11b* activation. However, a small but significant percentage of cells still failed to activate the enhancer-disrupted allele, and instead persisted in a monoallelic state with only expression of the Bcl11b-mCherry allele (Figures 3B-C, and S3-S4). Monoallelic cells were found in memory as well as naïve T-cell subsets (Figure S3), implying that these monoallelically expressing cells are capable of immune responses, as expected from the normal phenotype of Bcl11b knockout heterozygotes. As shown in fetal liver chimeras, generation and persistence of Bcl11b-mCherry monoallelic cells due to the mutant *Bcl11b*^YFPΔEnh^ allele were determined cell intrinsically (Figure S4). However, from flow cytometric profiles, cells that turned on the disrupted allele expressed it at normal levels, suggesting that the enhancer mutation reduced the stochastic rate of *Bcl11b* activation, but not its expression level once activated.

**Figure 3:**
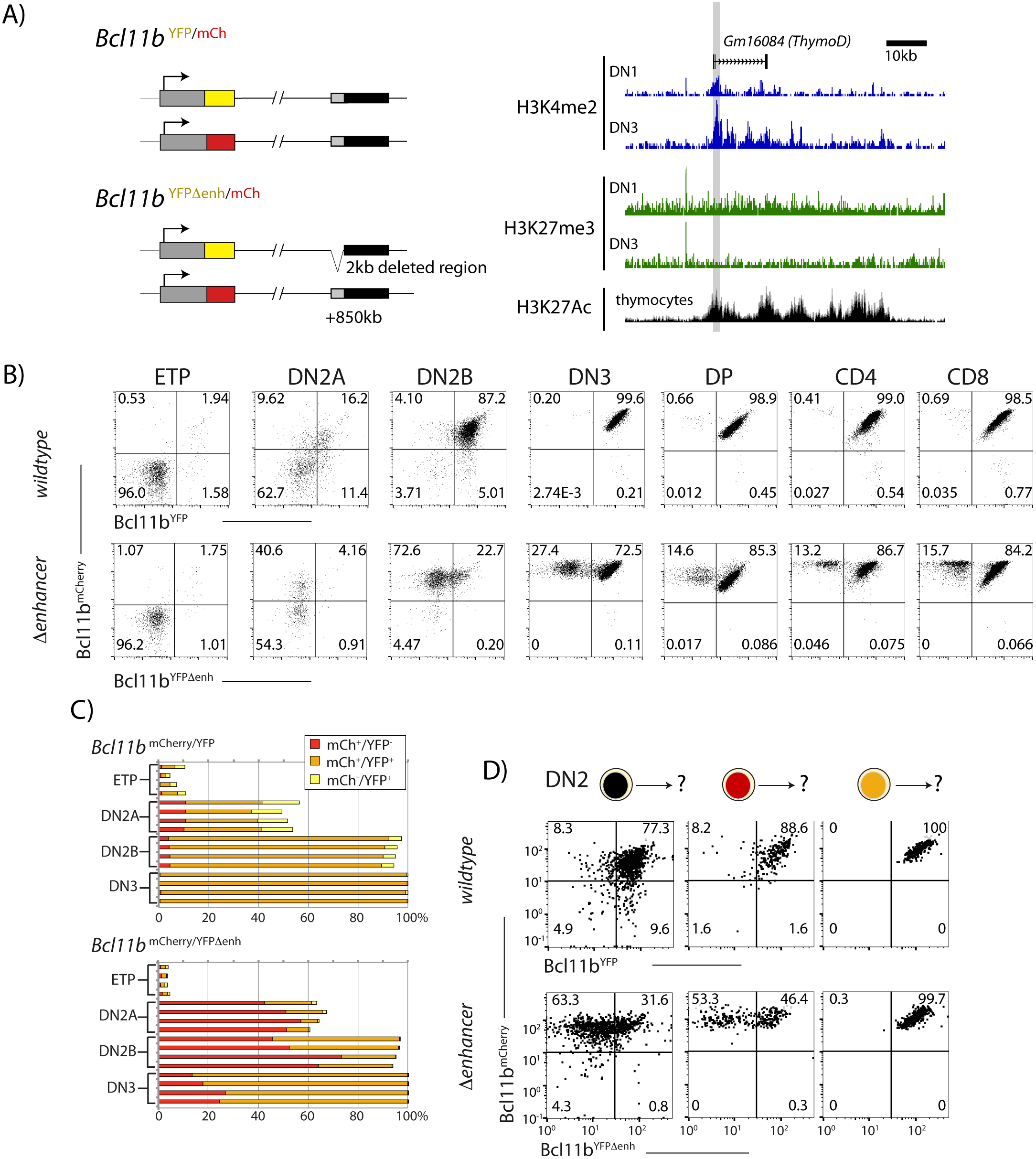
A distal enhancer region controls *Bcl11b* activation probability. A) Schematic of normal and enhancer-deleted two-color *Bcl11b* reporter strains (left). Genome browser plots (right), showing +850kb enhancer of *Bcl11b*, showing distributions of histone marks (H3K4me2, H3K27me, and H3K27Ac) and an associated LncRNA (Isoda et al., 2017). Orientation is with transcription from left to right (reversed relative to genome numbering). Gray shaded area indicates the enhancer region deleted using gene targeting (removed region: chr12:108,396,825-108,398,672, mm9). B) Flow cytometry plots show Bcl11b-mCh versus Bcl11b-YFP levels in developing T-cell populations from dual *Bcl11b* reporter mice, either with an intact YFP enhancer (top), or a disrupted YFP enhancer (bottom). Results are representative of 2 independent experiments. C) Bar graphs showing the percentages of cells in early thymic populations with mono- and biallelic expression of wildtype mCherry and wildtype versus mutant YFP in wildtype *Bcl11b*^YFP/mCh^ and *Bcl11b*^YFPΔEnh/mCh^ dual reporter mice, demonstrating the loss of mutant YFP allele expression relative to the wildtype mCherry allele in the same cells. Each bar shows results from one mouse; n = 4 mice of each strain are shown. D) DN2 progenitors were sorted for different *Bcl11b* allelic activation states as indicated, cultured on OP9-DL1 monolayers for 4 days, and analyzed using flow cytometry. Flow plots show Bcl11b-mCh versus Bcl11b-YFP levels of cells generated from precursors with a normal (top) or disrupted (bottom) YFP enhancer, showing defective YFP up-regulation from the mutant relative to the wildtype alleles. Enhancer disruption reduces the probability of switch-like *Bcl11b* activation, but does not affect expression levels after activation. Results are representative of 2 independent experiments. See also Figures S1, and S3 – S5.

To directly test this hypothesis, we measured Bcl11b-YFP activation with or without enhancer disruption, by sorting DN2 progenitors with zero or one allele activated, culturing on OP9-DL1 feeders, and analyzing activation dynamics of both alleles using flow cytometry. Consistently, enhancer disruption greatly reduced the fraction of cells that turned on Bcl11b-YFP, but did not perturb its expression level in cells that already successfully activated it (Figure 3D). Neither the wildtype nor the enhancer-disrupted allele reverted to silence after being activated. These results show that the deleted region within the distal *Bcl11b* super-enhancer works selectively, in *cis*, to accelerate the irreversible stochastic switch of the *Bcl11b* locus from an inactive to an active state.

The activation of the enhancer-disrupted *Bcl11b* allele observed in many DN2B and later cells suggests that there are other *cis*-regulatory elements on the *Bcl11b* locus that can also promote stochastic locus activation. The extended intergenic gene desert between *Bcl11b* and the next gene, *Vrk1*, is rich in potential regulatory elements that could compensate for the loss of the deleted enhancer element in the cells activating the YFP allele (Hu et al., 2018). Alternatively, the intact enhancer at the mCherry-tagged locus in the same cell could activate the enhancer-deleted Bcl11b locus in *trans*, but this was ruled out when we bred mice with the enhancer deletion to homozygosity (Bcl11b^YFPΔEnh/YFPΔEnh^). Progenitors from these mice were still able to turn on *Bcl11b* and to undergo T-cell development to CD4, CD8 double positive (DP) and single positive (SP) cells, and all the cells in these populations had normal levels of *Bcl11b* expression (Figure S5). Thus, the enhancer we identified works together with other regulatory elements specifically to control *Bcl11b* activation timing.

### A parallel trans-acting step enables expression from an activated Bcl11b locus

The known transcriptional regulators of *Bcl11b*—TCF-1, Gata3, Notch1 and Runx1—reach full expression prior to entering the DN2 stage, suggesting they are not limiting for *Bcl11b* activation in DN2 cells. The data presented above shows that *cis*-acting mechanisms substantially slow activation at individual alleles. However, additional *trans*-acting factors or post-translational changes in these factors could still limit the kinetics of *Bcl11b* activation, working either upstream of the *cis*-opening mechanism or as a separate, independent requirement. To gain insight into whether such *trans*-acting inputs are necessary to explain the observed dynamics, and how they could act together with *cis*-acting step, we developed a set of minimal models requiring the *cis-*activating step either alone or together with an additional *trans*-acting step (see Mathematical Appendix for model details).

In the simplest “*cis* only” model, we assume that only the *cis-*activation step is required for Bcl11b activation in DN2 stage, with all required *trans*-acting steps having occurred prior to the ETP-DN2 transition (Figure 4A, left). Because *cis-*activation is controlled at each allele by a single rate constant, this model predicts a lag between the appearance of monoallelic cells, which require one *cis-*activation event, and the appearance of biallelic cells, which require two independent events (Mathematical Appendix). By contrast, in experiments, some biallelic cells accumulated immediately, without a substantial lag relative to monoallelic ones, resulting in a poor fit of the data to the *cis-*only model (Figure 4A). These results rule out the simplest *cis*-only model, and suggest that additional *trans* events may still limit *Bcl11b* expression at the DN2 stage.

**Figure 4:**
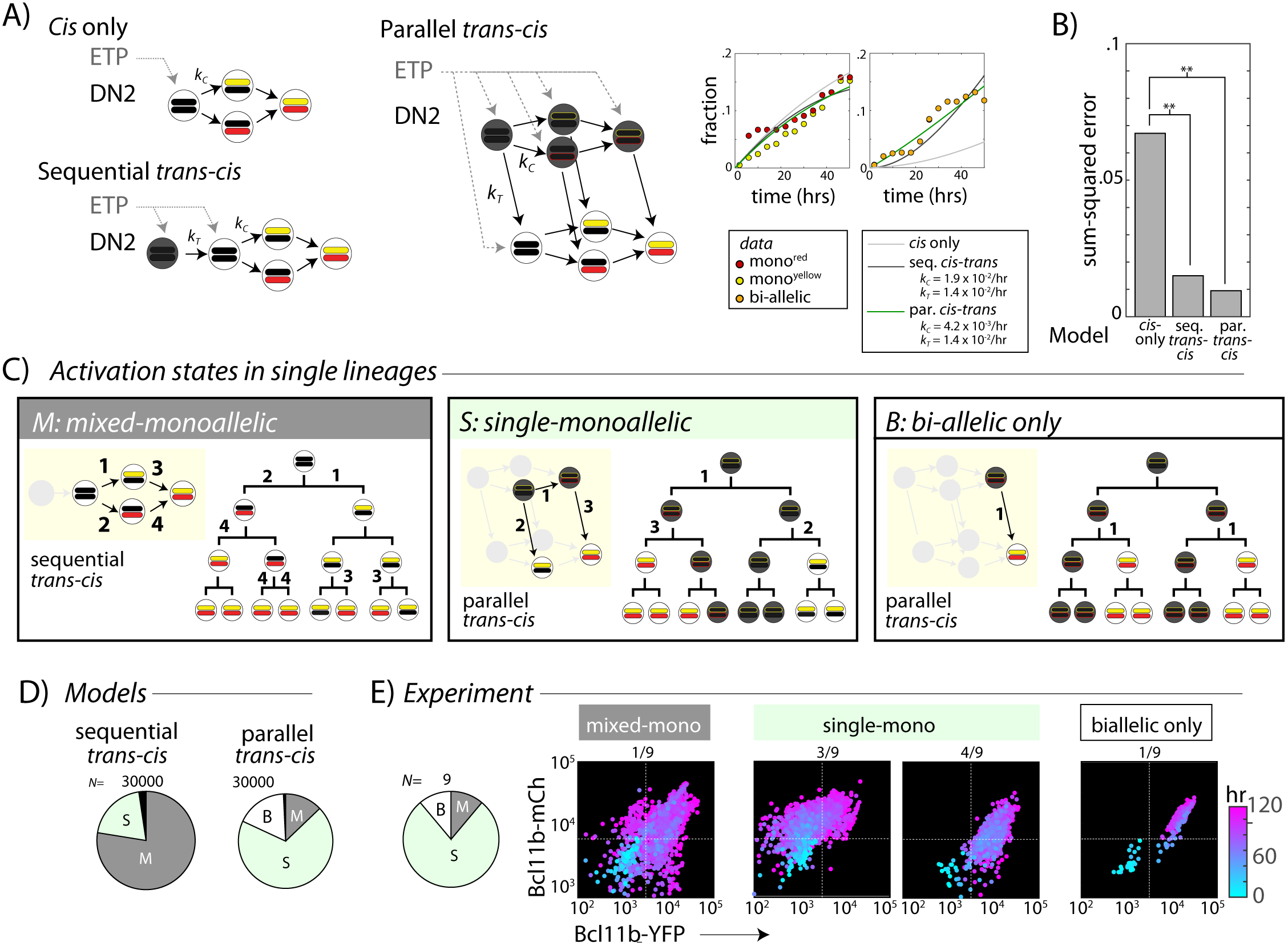
A *trans*-acting step, occurring in parallel with the *cis*-acting step, provides an additional input for *Bcl11b* activation. A) Candidate models for *Bcl11b* activation from the DN2 stage, involving a single *cis*-acting switch (top left), sequential *trans*-, then *cis*-acting switches (bottom left), and parallel, independent *trans*- and *cis*- acting switches (right). Plots show best fits of different models to the time evolution of *Bcl11b* allelic activation states, observed by timelapse imaging (Figure 2), with best fit rate constants indicated in legend. B) Bar charts show sum-squared errors for model fits, showing that both sequential and parallel *trans-cis* models fit the data significantly better than the *cis*-only model (F-test, *F*=17.4, p=0.0055, sequential vs. *cis*-only model; *F*=8.5, p=0.007, parallel vs. *cis*-only model). C) Three possible classes of *Bcl11b* activation states observable from clonal lineage data. Lineage trees and transition diagrams show examples of simulated lineages that fall into the indicated classes. D) Pie charts show expected distribution of allelic activation states predicted for clonal lineages of non-expressing progenitors in either the sequential (left) or the parallel (right) *trans-cis* model, obtained from *N*=30,000 simulations, using parameters derived from bulk fitting (see Mathematical Appendix). E) Pie chart (left) shows observed distribution of activation states observed across an entire imaging time course. Colored scatterplots (right) shows Bcl11b-mCh versus Bcl11b-YFP levels of single cell lineages, falling into the indicated categories. Clones were scored according to observable fluorescence across an entire developmental trajectory, from 0h (cyan) to 120 h (purple). The observed frequency of clones with ‘single-monoallelic’ expression of *Bcl11b* (7/9=77%) is significantly different than that predicted for the sequential *trans-cis* Model (20.4%, **-p<0.001, χ^2^ = 14.5, d.f. = 1), but not significantly different from that predicted for the parallel *trans-cis* Model (68.9%, χ^2^ = 0.1, d..f=1, n.s.). Results are representative of three independent experiments. See Figure S6 for data for independent replicate experiments.

We next considered two models in which *trans*-acting events affect *Bcl11b* activation. In the “sequential *trans-cis*” model, a *trans* step must occur prior to the *cis*-activation step (Figure 4A). This *trans* step could represent activation of a factor or epigenetic regulator that is necessary for *cis*-activation. In the “parallel *trans-cis*” model, both *cis* and *trans* steps are similarly necessary, but can occur in either order (Figure 4A). In this case, the *trans* step could represent activation of a factor that drives *Bcl11b* transcription, but only from a *cis*-activated locus. While our models only consider the DN2 stage, we note that they allow for some events to occur prior to the ETP-DN2A transition (Fig. 4A, gray dotted arrows). When the *trans*-acting step is rate limiting, both of these models reduce biallelic lag by allowing the two alleles to turn on in relatively quick succession (in either model) or simultaneously (in the parallel model). For this reason, both the sequential and parallel *trans-cis* models reduced the lag prior to accumulation of biallelic cells, and hence fit the data significantly better than the “*cis* only” model (Figures 4B, p < 0.01 for both models).

While the sequential and parallel models show similar bulk behavior, they make divergent predictions about the distributions of mono- and bi-allelic expression states within clonal lineages. For example, in the sequential model, silent progenitors are equally likely to activate one or the other *Bcl11b* allele, and are thus more likely to show mono-allelic expression from both alleles in single clones (Fig. 4C, “mixed monoallelic”). In contrast, in the parallel model, non-expressing progenitors could have one *cis*-activated but unexpressed *Bcl11b* allele due to absence of the *trans* step. Clonal descendants of such cells would be predisposed to show mono-allelic expression from the same allele before activating the second (Fig. 4C, “single monoallelic”). Therefore, to discriminate between sequential and parallel activation models, we used Monte-Carlo methods to simulate the dynamics of *Bcl11b* activation in all descendants of a single starting cell over four generations for each of the two models (Mathematical Appendix), using the parameters that gave the best fits to the global time course data in Fig. 4A. Altogether, we generated and analyzed N=30,000 clonal lineages for each model.

As intuitively expected, the sequential *trans-cis* model predominantly generated ‘mixed monoallelic’ clones containing cells with mono-allelic expression of both alleles, with or without biallelically expressing cells (Figure 4C-D, “mixed monoallelic”). These distributions reflect the most likely event trajectory in the sequential model, in which independent, unsynchronized *cis*-activation events occur at each *Bcl11b* locus in different cells from a single ancestor. Within a cohort of clonal descendants competent to activate the *cis-*step, the first-activated allele choice occurs independently in each cell, generating multiple paths towards biallelic *Bcl11b* activation within a single clone. By contrast, the parallel model generated a much smaller fraction of such “mixed monoallelic” clones, and predominantly generated clones in which monoallelic expression was restricted to the same allele across most cells (Figure 4C-D, “single monoallelic”). This intra-clonal bias arises when the *cis*-acting step at one locus precedes the *trans*-step, forcing still-inactivated DN2A precursors to preferentially activate that locus once the *trans*-acting event occurs (Fig. 4C). Because the rate of *cis*-activation is low (τ_c_ ~ 4-6 days), individual cells within a clone can monoallelically activate the same locus prior to full biallelic expression. Moreover, the parallel but not the sequential *trans-cis* model gave rise to a small fraction of clones that showed only bi-allelic expression (Fig. 4C-D, “bi-allelic only”), reflecting the activation of the *trans*-limiting step in cells that had already undergone *cis*-activation of both *Bcl11b* copies.

To discriminate experimentally between these two models, we quantified the distribution of *Bcl11b* allelic activation states generated in clonal lineages from progenitors starting with no *Bcl11b* activation, observed by timelapse microscopy as described above (Fig. 2). Within a clone, we most frequently observed monoallelic expression from only one specific allele, with or without biallelically expressing cells (Fig. 4E “single monoallelic”, light green, 7/9 clones, Fig. S6; similar results observed over three independent experiments), but only rarely observed monoallelic expression from both loci within the same clone (Fig. 4E, “mixed mono-allelic”, gray, 1/9 clones). The observed percentage of “single monoallelic” expressing clones (7/9 = 77%) was significantly greater than that expected from the sequential *trans-cis* model (20.4%, p < 0.005). Moreover, in one clone, we observed concurrent activation of both alleles (Fig. 4E), a behavior that would have been exceedingly rare in a sequential model (none observed in 30,000 simulations). Together, these results suggest that a *trans*-acting step, acting in parallel with the *cis-*acting step, controls *Bcl11b* expression.

### Notch signaling controls the parallel trans-acting step in Bcl11b activation

Notch signaling drives T-cell fate commitment and provides an important input for *Bcl11b* expression. While not required to maintain *Bcl11b* expression in committed cells, it acts earlier to enhance the probability of all-or-none *Bcl11b* expression at the DN2 stage and stabilize *Bcl11b* expression shortly after activation, preventing the re-silencing that still can occur in a small fraction of newly expressing cells (Kueh et al., 2016). The Notch intracellular domain is diffusible in the nucleus, but could affect *Bcl11b* activation by modulating either the *cis* or *trans* step in the parallel model. For example, Notch signaling could activate a *trans* factor that drives *Bcl11b* transcription from a *cis*-activated locus. Alternatively, Notch could affect the rate of the *cis-*activation process, for instance by enhancing the activity of chromatin-remodeling enzymes on the *Bcl11b* promoter or enhancer (cf. Fig. 3).

To distinguish between these two potential roles of Notch, we analyzed the effects of Notch signaling withdrawal on *Bcl1b* allelic expression patterns and compared the results to predictions from the parallel *trans-cis* model. For these assays, we first sorted DN2 cells with no expression, mono-allelic expression or bi-allelic expression of *Bcl11b* as initial populations, and then cultured them either on OP9-DL1 or OP9-control feeders to maintain or remove Notch signaling, respectively. After four days, we analyzed the resulting Bcl11b allelic expression states using flow cytometry (Fig. 5A).

**Figure 5:**
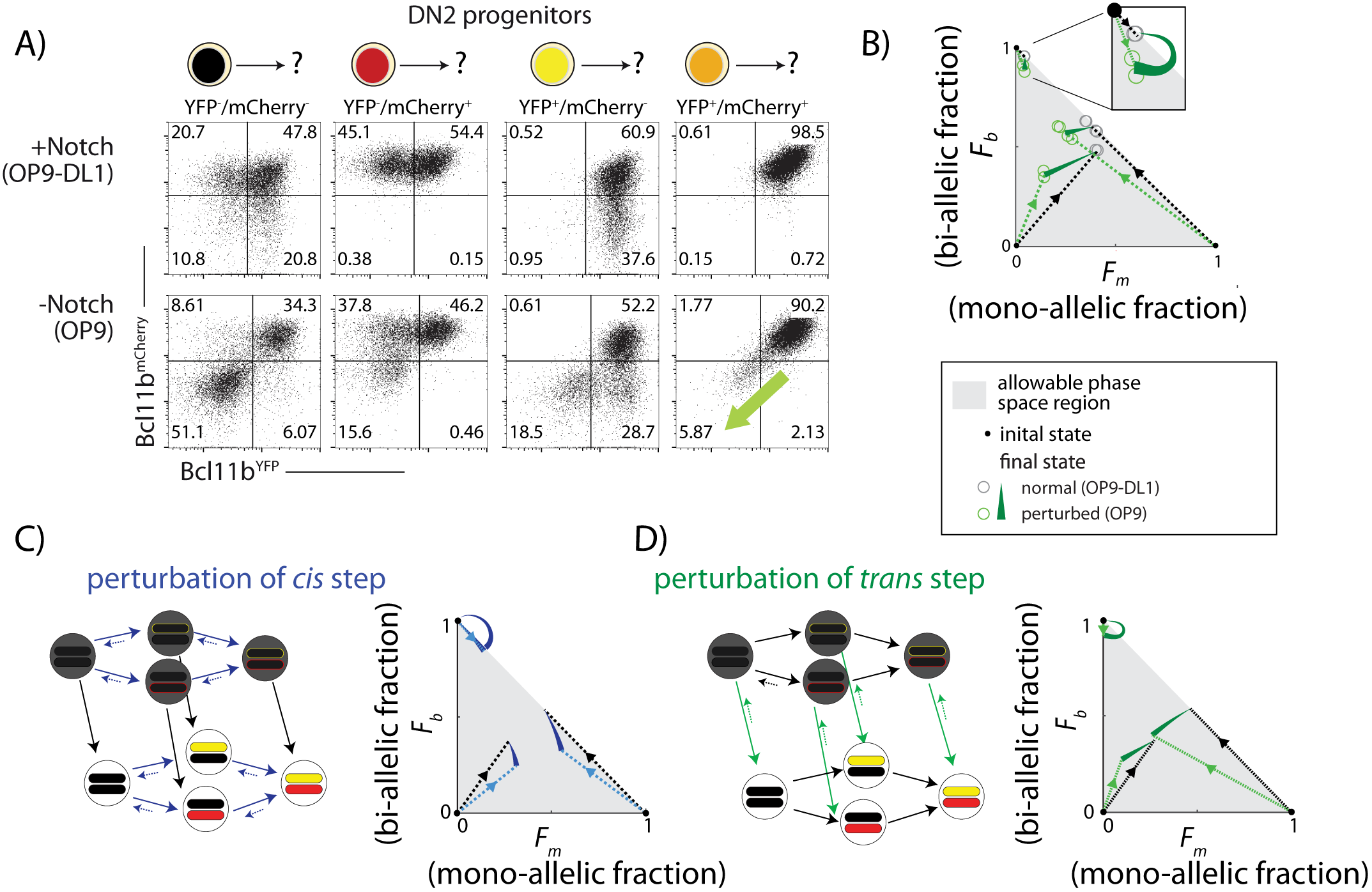
Notch signaling controls a parallel *trans*-acting step for *Bcl11b* activation. BM-derived DN2 progenitors with different *Bcl11b* allelic activation states were sorted, cultured on either OP9 (-Notch) or OP9-DL1 (+Notch) monolayers for four days, and analyzed using flow cytometry. A) Flow cytometry plots show Bcl11b-mCherry versus Bcl11b-YFP expression levels in analyzed cells. Percentages of non-expressing, monoallelic expressing (both YFP and mCherry) and biallelic expressing cells were used to calculate the locations in the phase space. Note that when Notch signaling is withdrawn from biallelically expressing cells, they downregulate both alleles coordinately (green shaded arrow). B) Phase space diagrams experimentally obtained from analysis of flow cytometry data. Points in phase space represent the average of 2-4 replicate data points in a single experiment (hollow circles). Inset shows final activation states of bi-allelic starting progenitors upon Notch withdrawal. Results shown are representative of 3 independent experiments. C)-D) Predicted phase space diagrams for fraction of biallelic expressing cells (*F_b_*) against the fraction of monoallelic expressing cells (*F_m_*, YFP+ and mCh+ combined), for either the sequential *trans-cis* activation model (C), or the parallel *trans-cis* model (see Mathematical Appendix for details). Black (colored) dotted lines connect initial state to the normal (perturbed) final state. Note that actual developmental trajectories may be curved (not shown). Arrows show predicted shifts in final state due to the indicated perturbations. Note that perturbations affect both the rates and reversibility of the indicated reactions. See also Figure S7.

DL1 removal caused distinct shifts in the distributions of final *Bcl11b* allelic expression states across each of the starting cell states. For progenitors with no initial *Bcl11b* expression, DL1 withdrawal decreased the total fraction of cells that subsequently expressed *Bcl11b* from either allele (from 0.9 to 0.5, sum of mono-allelic and bi-allelic expressing cells, Fig. 5A), consistent with previous results (Kueh et al., 2016). DL1 withdrawal differentially affected the mono-allelic expressing population, such that the ratio of mono-allelic to bi-allelic expressing cells fell from ~0.8 to 0.4 (Fig. 5A). In progenitors starting with mono-allelic *Bcl11b* expression, DL1 removal inhibited expression of the initially silent allele, and led to inactivation of the initially expressing allele in a small fraction of cells (Fig. 5) (Kueh et al., 2016). As with the non-expressing progenitors, it also reduced the ratio of mono-allelic to bi-allelic expressing cells (Figures 5A, 5B). Finally, in bi-allelic *Bcl11b* progenitors, most cells maintained expression despite Notch removal as expected (Kueh et al., 2016), but a small fraction (~0.06) lost expression of both *Bcl11b* alleles entirely, reverting directly from the bi-allelic to a non-expressing state (Fig. 5A, diagonal arrows). The distribution of non-expressing, monoallelic, and biallelic expressing cell states in the population in any condition can be represented as a point in a triangular region of allowed states in a single diagram (Fig. 5B), providing a visual summary of the effect of Notch withdrawal on the population (Figure 5B).

We compared these effects of experimental Notch withdrawal with predicted effects of a step-like perturbation in either the *cis* or the *trans*-acting steps (see Mathematical Appendix). In order to account for reversibility in Bcl11b activation observed upon DL1 removal (Fig. S7), each perturbation was assumed to both decrease the rate of the forward (*cis* or *trans*) step and increase the rate of a reverse step. Simulations of the resulting models generated distributions of *Bcl11b* allelic activation states from non-expressing, monoallelic, and biallelic starting populations, with no perturbation, or with perturbation of the *cis* or *trans*-acting steps.

Perturbation of the *cis*-acting step decreased the total fraction of cells expressing *Bcl11b* from all initial cell populations, as expected. However, in contrast to experimental observations, this simulated perturbation increased, rather than decreased, the ratio of mono-allelic expressing cells to bi-allelic expressing cells (Fig. 5C). It also caused biallelic expressing cells to sequentially turn off Bcl11b one allele at a time, rather than simultaneously as observed experimentally. These results suggest that perturbation of the *cis-*acting step does not account for the observed effects of Notch withdrawal (Fig. 5B,C).

By contrast, perturbation of the *trans*-acting step in the model produced effects resembling Notch withdrawal. First, it decreased the ratio of mono-allelic to bi-allelic expressing cells for starting progenitors with no or mono-allelic *Bcl11b* expression. Second, it led to direct reversion of bi-allelic expressing progenitors to a non-expressing state, without passing through mono-allelic intermediates (Fig. 5D, green arrows). Concurrent inactivation of both alleles is difficult to reconcile with Notch affecting independent (*cis*) effects at each allele, but is expected in response to removal of a *trans*-acting factor required for maintaining expression (Figure 5B). Additionally, we note that these observations were also inconsistent with Notch controlling a necessary *trans*-acting step occurring strictly prior to *cis*-activation, as postulated by the sequential *trans-cis* model (Figure S7). In this case, progenitors that express one or both *Bcl11b* alleles would no longer be affected by Notch withdrawal, inconsistent with our observations (Figure 5B). Taken together, these results strongly suggest that a separate Notch-dependent *trans*-acting event, occurring in parallel with Bcl11b locus activation, is necessary for Bcl11b activation and T-cell lineage commitment.

### Bcl11b activation can only occur over a limited developmental window

Given the finite rate of *cis-* and *trans-*activation steps, all cells would be expected to eventually activate both *Bcl11b* copies. However, a small fraction of cells were consistently found to express *Bcl11b* monoallelically in thymic and peripheral T cell subsets (Figures 3B-C, S3, and S4). This result suggested that cells might lose competence to activate any still-silent *Bcl11b* locus as they develop. To test this hypothesis, we sorted monoallelically expressing cells from different developmental stages, cultured them *in vitro* on OP9-DL1 monolayers for four days, and analyzed expression of both *Bcl11b* alleles (Fig. 6A). The already-active copy retained active expression throughout the assay, as expected. However, the frequency of activation of the initial silent *Bcl11b* allele varied strongly with developmental stage. Activation occurred efficiently at the DN2 stage (DN2A and DN2B combined) but dropped sharply as cells progressed to DN3 (~80% versus ~15% activated after four days, Figure 6A), and dropped even further at the double positive (DP) and CD4 single positive stages (~1.5% and 2.4%, respectively, Fig. 6A). Equivalent results were obtained regardless of whether the experiment started with active YFP and mCherry alleles (Fig. 6A). These results indicate that *cis-*activation of *Bcl11b* predominantly occurs during DN2 and DN3 stages.

**Figure 6:**
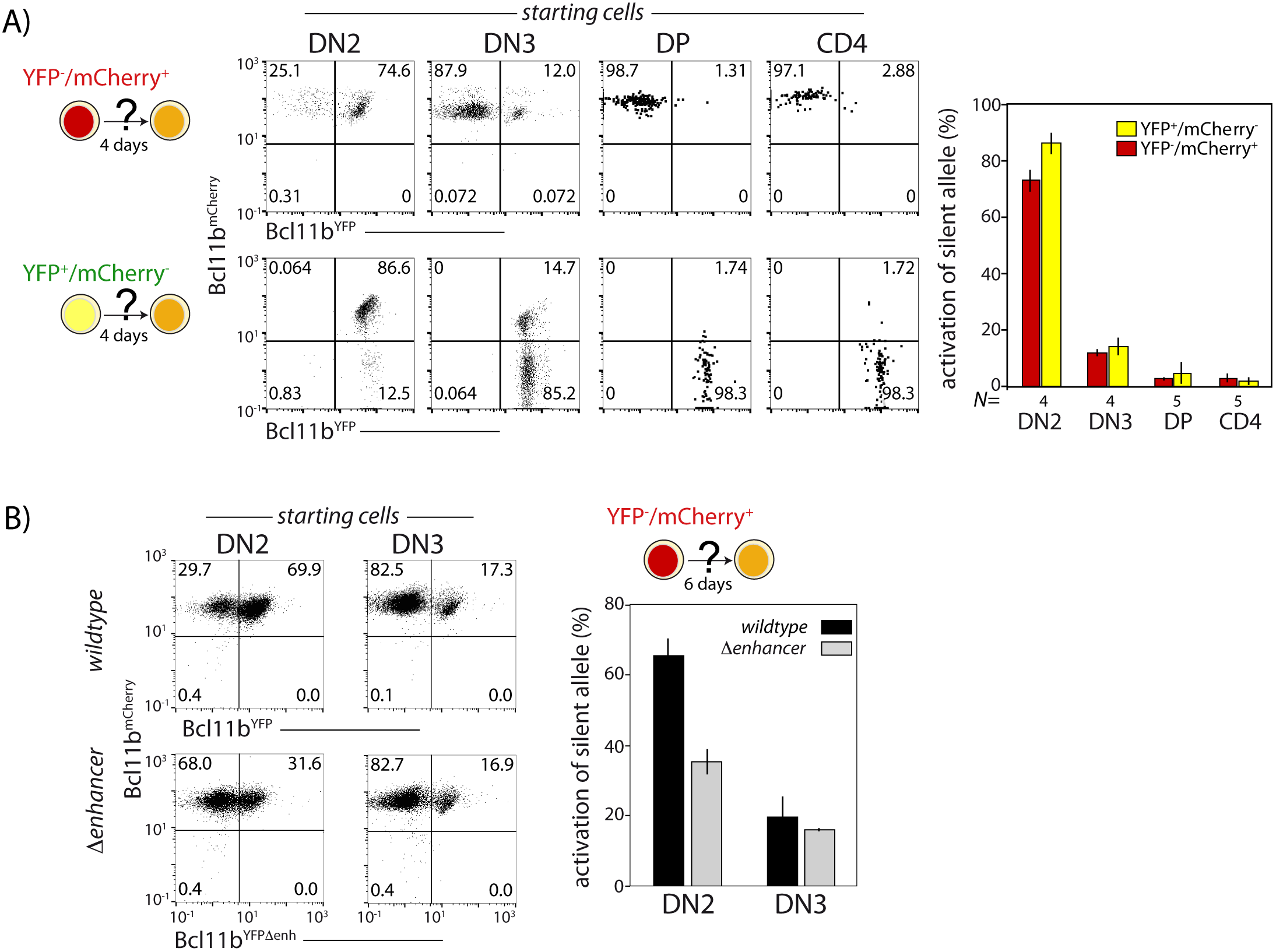
Probabilistic *Bcl11b* activation occurs within a limited developmental time window. Cells expressing only one *Bcl11b* allele at the indicated stages were sorted from thymocytes, cultured for 4d on OP9-DL1 monolayers, and analyzed for activation of the initially inactive *Bcl11b* allele using flow cytometry. A) Flow plots (left) show Bcl11b-mCh versus Bcl11b-YFP expression levels for descendants of cells that had monoallelic expression at the indicated stages of development; bar charts (right) show the fraction of progenitors from different stages that activate the silent *Bcl11b* allele upon culture. Data represent mean and standard deviation of 4-5 replicates, derived from 2 independent experiments. The competence to activate the silent *Bcl11b* allele decreases upon progression to the DN3 stage and beyond. B) Flow plots (left) show Bcl11b-mCh versus Bcl11b-YFP expression levels for DN2 or DN3 progenitors with either an intact YFP allele enhancer (top) or a disrupted YFP allele enhancer (bottom). Bar chart (right) shows the fraction of cells activating the silent *Bcl11b* allele upon re-culture. Data show that enhancer disruption reduces the *Bcl11b* activation advantage in DN2 cells as compared to DN3 cells. Data represent mean and standard deviation of 3 replicates from 2 independent experiments.

**Figure 7:**
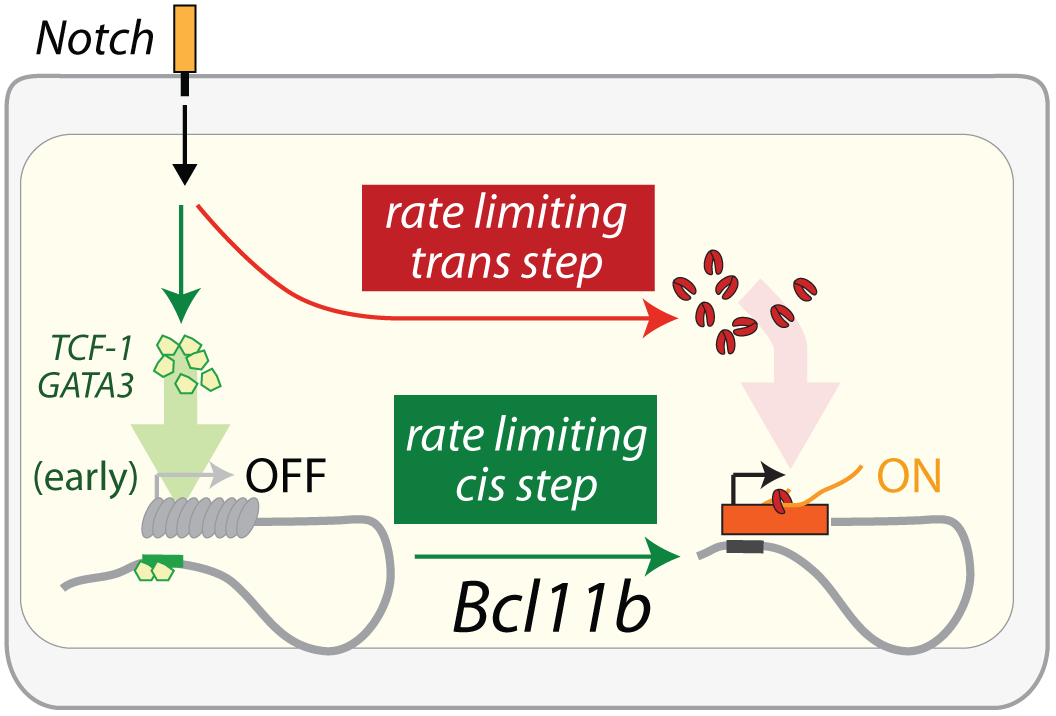
Model of *Bcl11b* regulation by parallel *cis* and *trans* limiting steps. *Bcl11b* activation requires two rate-limiting steps: a switch of the *Bcl11b* locus from an inactive to active epigenetic state, and the activation of a *trans* factor is necessary for transcription of *Bcl11b* from an activated locus. Notch signaling activates TCF-1 and GATA3 in early thymic progenitors (García-Ojeda et al., 2013; Scripture-Adams et al., 2014; Weber et al.), and these two factors may act on the identified distal enhancer to control the rate-limiting *cis* step on the *Bcl11b* locus (green). In parallel, Notch promotes the activation of a *trans* factor (red) that is necessary for transcription from a *cis*-activated *Bcl11b* locus. The *cis* and *trans* limiting steps together control the dynamics of Bcl11b expression and T-cell lineage commitment.

This DN2-stage preference for *Bcl11b* activation competence could arise from stage-specific activity of the identified distal enhancer. To test this hypothesis, we compared the activation kinetics of intact and enhancer-disrupted YFP alleles in sorted progenitors expressing only the *Bcl11b* mCherry allele. When the input cells were DN2 cells, the enhancer-disrupted YFP allele showed markedly less activation over the next four days than the intact YFP allele (70% versus 32%, Figure 6B). However, using input cells sorted at the DN3 stage, no differences in activation propensity were observed, with both wildtype and disrupted enhancer alleles showing the same attenuated degree of activation (~17%). These results suggest that the *Bcl11b* enhancer works specifically to enhance *cis*-activation of *Bcl11b* at the DN2 stage.

## Discussion

Stochastic epigenetic control switches have been described in yeasts, plants, and, more recently, constructed in synthetic systems (Berry et al., 2017; Bintu et al., 2016; Hathaway et al., 2012; Keung et al., 2014; Xu et al., 2006), yet their roles in controlling fate decisions in vertebrate developmental systems are not well understood. Specifically, it is not clear when epigenetic states simply respond passively to ‘upstream’ developmental changes in transcription factor activity, and when they actively impose distinct temporal constraints on transcription factor effects. By separately following the two chromosomal copies of *Bcl11b* in single cells, we found that the decision to turn on *Bcl11b*, and the ensuing transition to T-cell fate, involves a stochastic, irreversible rate-limiting *cis-*activation step that occurs on the each chromosomal allele of the *Bcl11b* gene itself. The *cis*-acting step occurs at a low enough rate (*k_C_* = (4.2 ± 3.3)×10^−3^/hr, Figure 4A) to generate numerous monoallelically expressing cells as intermediates, and is stable enough to propagate the same monoallelic activation state through multiple rounds of cell division in individual clones. In particular, by generating delays of multiple days and cell generations prior to differentiation, the *cis-*acting switch also indirectly controls the overall degree of proliferation of the progenitor pool. These results thus demonstrate that stochastic, epigenetic events on individual gene loci can fundamentally limit the timing and outcome of mammalian cell fate decisions, as well as the population structure of the resulting differentiated population.

Slow, stochastic *Bcl11b* activation is controlled by an enhancer far downstream from the *Bcl11b* promoter, on the opposite end of the same topologically associated domain. Multiple known epigenetic changes that occur on the *Bcl11b* locus could participate in the processes whose dynamics we have measured here. The distal enhancer could recruit chromatin regulators that clear repressive chromatin modifications from the *Bcl11b* locus. In its silent state, the *Bcl11b* promoter and gene body are covered by DNA methylation and histone H3K27me3 modifications (Hu et al., 2018; Ji et al., 2010; Zhang et al., 2012). Chromatin regulators recruited by the enhancer could disrupt repressive modifications in their vicinity, catalyzing a phase transition that results in cooperative, all-or-none removal of repressive marks on the entire gene locus (Larson et al., 2017; Strom et al., 2017). As another possibility, the distal enhancer could recruit *trans*-factors that facilitate its T-lineage-specific looping with the *Bcl11b* promoter (Li et al., 2013). As *Bcl11b* turns on, its promoter establishes new contacts with the distal enhancer, resulting in *de novo* formation of an altered topological associated domain, with boundaries defined by these two elements (Hu et al., 2018; Isoda et al., 2017). *Trans-* regulators of DNA loop extrusion that associate with the distal enhancer, whose binding may be facilitated by non long-coding RNA transcription (Isoda et al., 2017), may stabilize these looping interactions (Fudenberg et al., 2016; Nasmyth, 2001; Riggs, 1990; Sanborn et al., 2015). The evidence for such epigenetic differences associated with the *Bcl11b* locus in T and non-T have been known for some time, but the functional impacts of *cis*-acting mechanisms on locus activation dynamics has been unknown until now. Ultimately, any of these mechanisms that are rate-limiting will have to account for the stochastic nature of *Bcl11b* locus activation, its exceptionally long activation time constant, and its all-or-none, irreversible nature, demonstrated here. Dissecting the molecular and biophysical basis of these striking emergent properties will be the subject of future investigation.

Our mathematical models, together with perturbation analysis, further show that *Bcl11b* expression also requires a separate Notch signal-dependent *trans*- event that is needed in parallel with *Bcl11b cis-*activation. Given the comparable slow rate constants for parallel *cis* and *trans* steps in our model, it is expected that a substantial fraction of cells would undergo the *cis*-acting step prior to *trans*-activation and observable *Bcl11b* expression. It is even possible that the *cis*-acting step could occur earlier, i.e. within the ETP stage or during the ETP-DN2a transition (Kueh et al., 2016). In fact, earlier *cis-*activation is consistent with previous results showing ETP stage-specific dependence on the transcription factors GATA-3 and TCF-1 in controlling *Bcl11b* locus activation (Kueh et al., 2016), as well as chromatin changes occurring at the *Bcl11b* distal enhancer already observed at the ETP stage (Isoda et al., 2017; Zhang et al., 2012).

Slow, stochastic epigenetic switches, like the one described here, may allow cells to tune the size and composition of differentiated tissues. By using *trans*-acting inputs that modulate activation probabilities, such epigenetic switches could translate differences in input duration to changes in the fraction of output cells activated (Bintu et al., 2016), a strategy that could enable tunable control of cellular proportions in a developing tissue or organ. Moreover, a striking aspect of this mechanism is its ability to generate populations of mature T cells that are mosaic in the status of their activation of the two *Bcl11b* alleles. Indeed, the differential distribution of monoallelically expressing cells that we see among distinct functional T-cell subsets suggests the potential of non-uniform allelic activity to alter function or selective fitness. The increased fraction of monoallelically expressing cells that appear when an enhancer complex is weakened is a strong phenotype at the single cell level that could be relevant to enhancer polymorphisms in natural populations, although its impact could easily be underestimated by more conventional gene expression analyses.

Here, we have illustrated a general approach that can reveal the dynamics of epigenetic control mechanisms, determine their prevalence in the genome, and elucidate their functional roles in multicellular organism development and function. Stochastic epigenetic switches, similar to the one uncovered here, may constitute fundamental building blocks of cell fate control circuits in mammalian cells. As cells transition from one developmental state to another, they undergo concerted transformations in the chemical modification states or physical conformations of many regulated genes. These changes could reflect more widespread roles for epigenetic mechanisms in controlling cell state transition timing.

## Acknowledgements

We thank M. Lerica Gutierrez Quiloan for mouse genotyping and maintenance; N. Verduzco and I. Soto for animal husbandry; R.A. Diamond, K. Beadle, and D. Perez for cell sorting. We also thank members of Kueh, Rothenberg and Elowitz labs for feedback, and T. Mitchison for valuable discussions. We also thank Sandy Nandagopal, Pulin Li, Zeba Wunderlich and Nick Pease for comments. This work was funded by an NIH K99/R00 Award (5R00HL119638), a Tietze Foundation Stem Cell Scientist Award, and a CRI/Irvington Postdoctoral Fellowship (to H.Y.K.); NIH grants R01AI095943, R01AI083514, and R01HL119102 (to E.V.R.), California Institute for Regenerative Medicine Bridges to Stem Cell Research (to K.K.H.N.); and the Louis A. Garfinkle Memorial Laboratory Fund, the Al Sherman Foundation, and the Albert Billings Ruddock Professorship (to E.V.R.).

---

**Figure S1:**
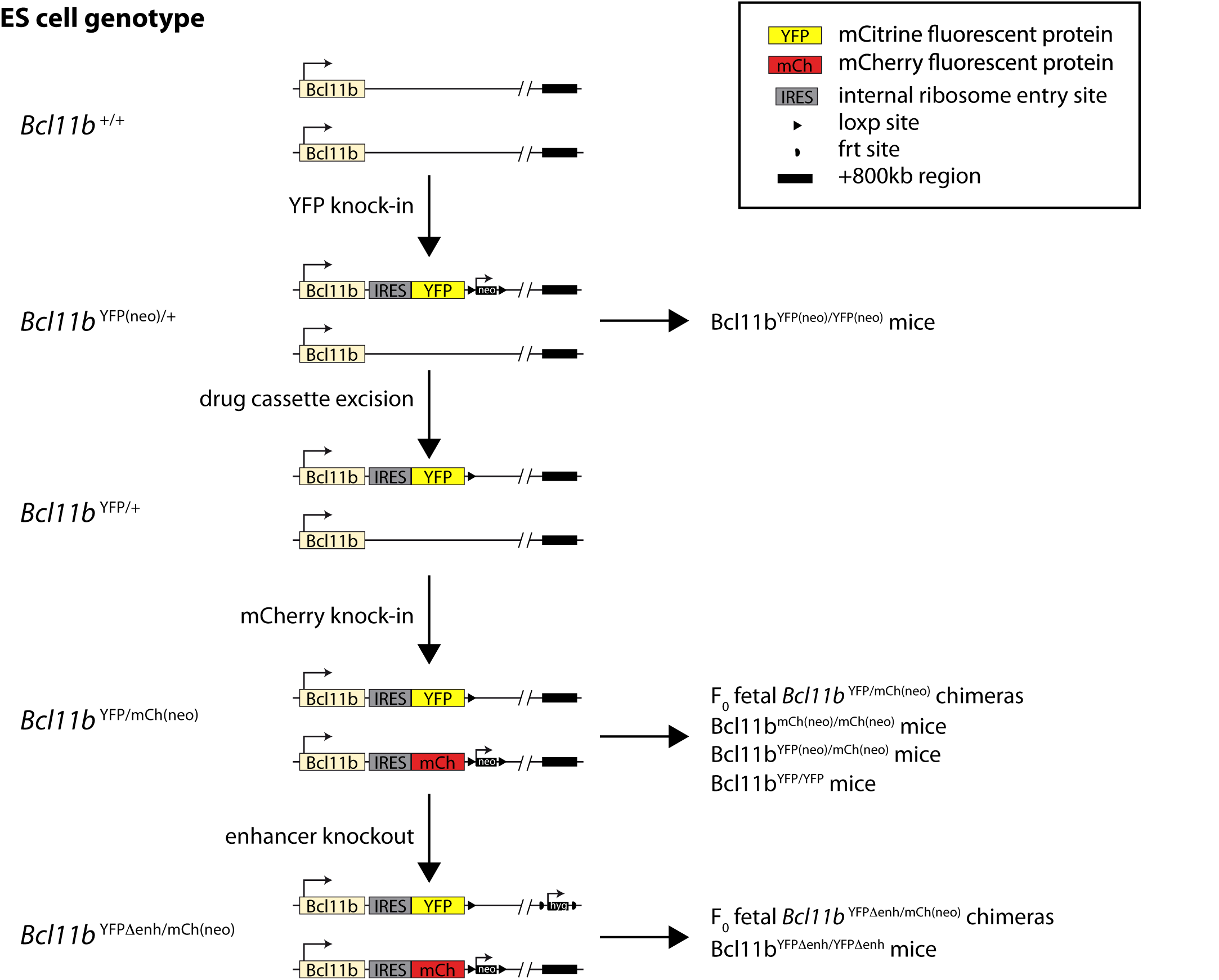
Experimental strategy for generating different *Bcl11b* reporter mouse strains, Related to Figures 1 and 3. The two *Bcl11b* loci were targeted in embryonic stem (ES) cells using homologous recombination, followed by drug selection using the indicated drug-resistance markers (vertical arrows). ES cells were then injected into blastocysts (horizontal arrows) to generate the indicated mice. These mice were subsequently bred to generate the appropriate two-color mice for experiments (see Methods). To generate the enhancer disrupted ES cells, the dual-color tagged ES line was retargeted with a deletion construct including a selectable hygromycin resistance gene (*hyg*). Cells with the correct insertion were then used to generate mice.

**Figure S2:**
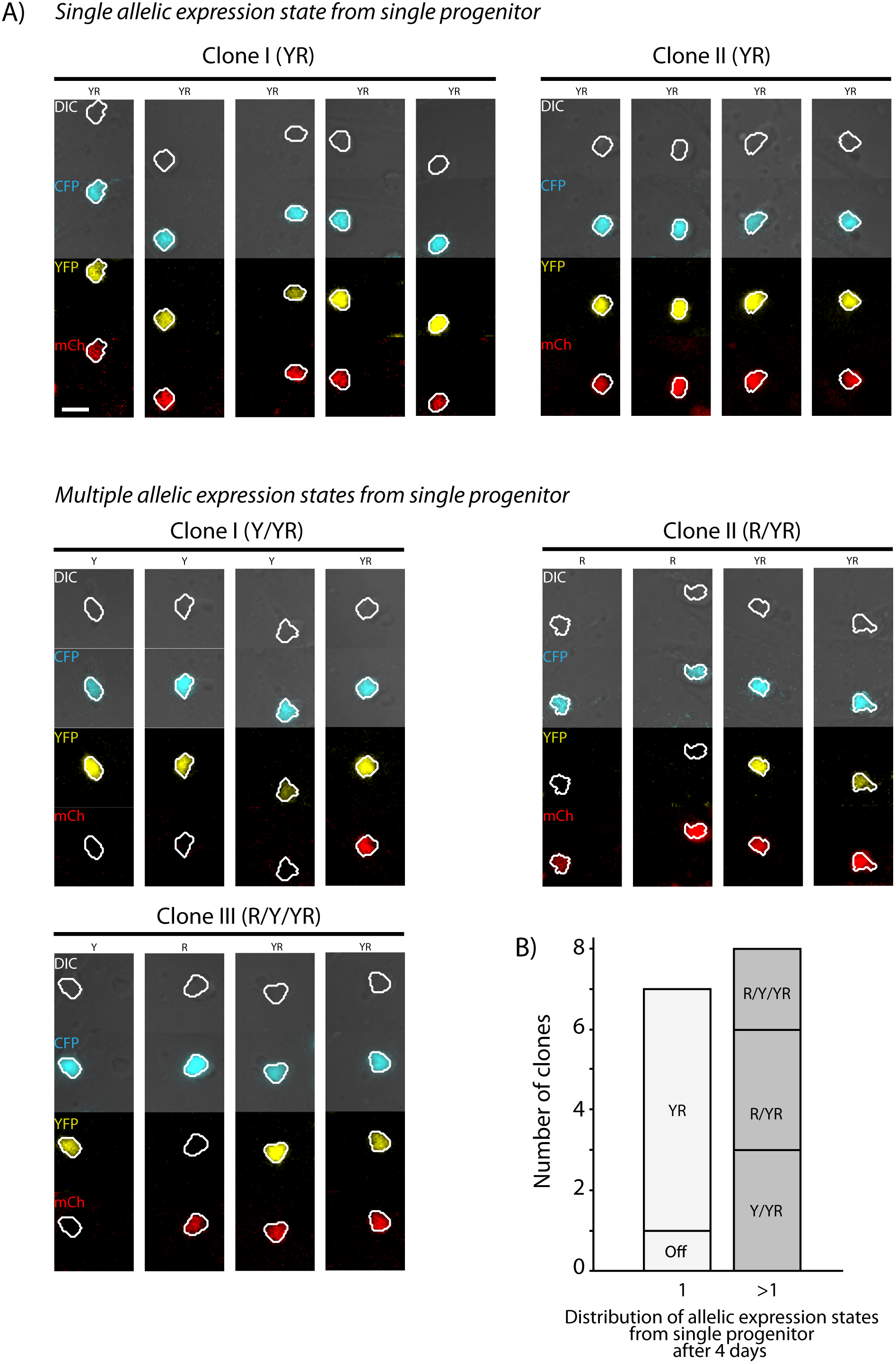
*Bcl11b* shows heterogeneity in locus activation within clonal progenitor lineages, Related to Figures 2 and 4. Bone marrow derived Bcl11b-YFP^-^mCherry^-^ DN2 progenitors were sorted, seeded on OP9-DL1 monolayers within PDMS micro-well arrays, and continuously observed using long-term timelapse imaging. Microwells seeded with single proliferating cell clones were identified, and *Bcl11b* activation states of descendants were then analyzed after four days. A) Timelapse images show DIC (gray), YFP (yellow) and mCherry (red) fluorescence of cells descended from single progenitors. Each row represents cells within a single microwell. Images were taken between 105 and 115 hours after onset of imaging. Scale bar represents 10 microns. B) Stacked bar chart shows the range of *Bcl11b* activation states observed for clonal descendants after four days. Left bar indicates clones where cells were all found in the same activation state; right bar gives clones with multiple activation states observed within a single clone.

**Figure S3:**
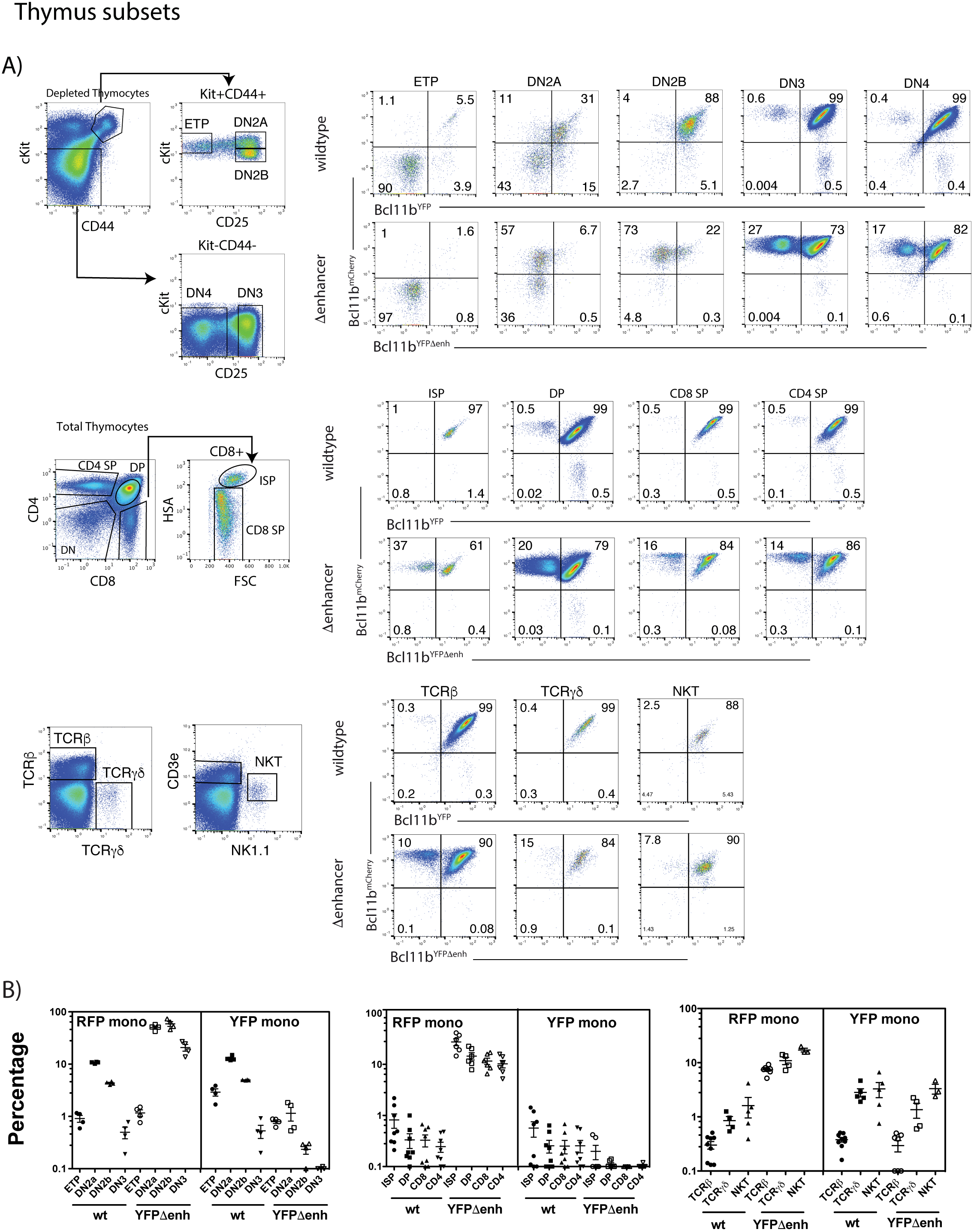
Levels of monoallelic *Bcl11b* expression in thymus subsets: monoallelic expression can persist throughout thymic development, Related to Figure 3. A) Representative flow cytometry plots showing gating strategies for thymic subsets and two-color Bcl11b expression in these populations from Bcl11b^YFP/mCh(neo)^ (wildtype) or Bcl11b^YFPΔEnh/mCh(neo)^ (Δenhancer) mice. DN subsets were enriched by magnetic bead depletion of mature thymic cells before staining and analysis. B) Percentages of cells expressing only mCherry (RFP mono) or YFP (YFP mono) in specific T cell populations from Bcl11b^YFP/mCh(neo)^ (wt) or Bcl11b^YFPΔEnh/mCh(neo)^ (YFPΔenh) mice. Each symbol represents results from an individual mouse (n=4 to 6 mice per group). This figure shows that although biallelic expression predominates, monoallelic expression of both YFP and mCherry wildtype alleles persist in some cells throughout intrathymic development. Furthermore, the YFPΔenh mutant dramatically increases the percentage of cells expressing only the mCherry (wildtype) allele due to failure to activate the mutant allele. However, the level of monoallelic expression seen decreases generally over development of CD4 and CD8 SP αβ T cells and is slightly higher among TCRγδ^+^ and NKT cells relative to conventional TCRβ^+^ cells, possibly consistent with additional selection events.

**Figure S4:**
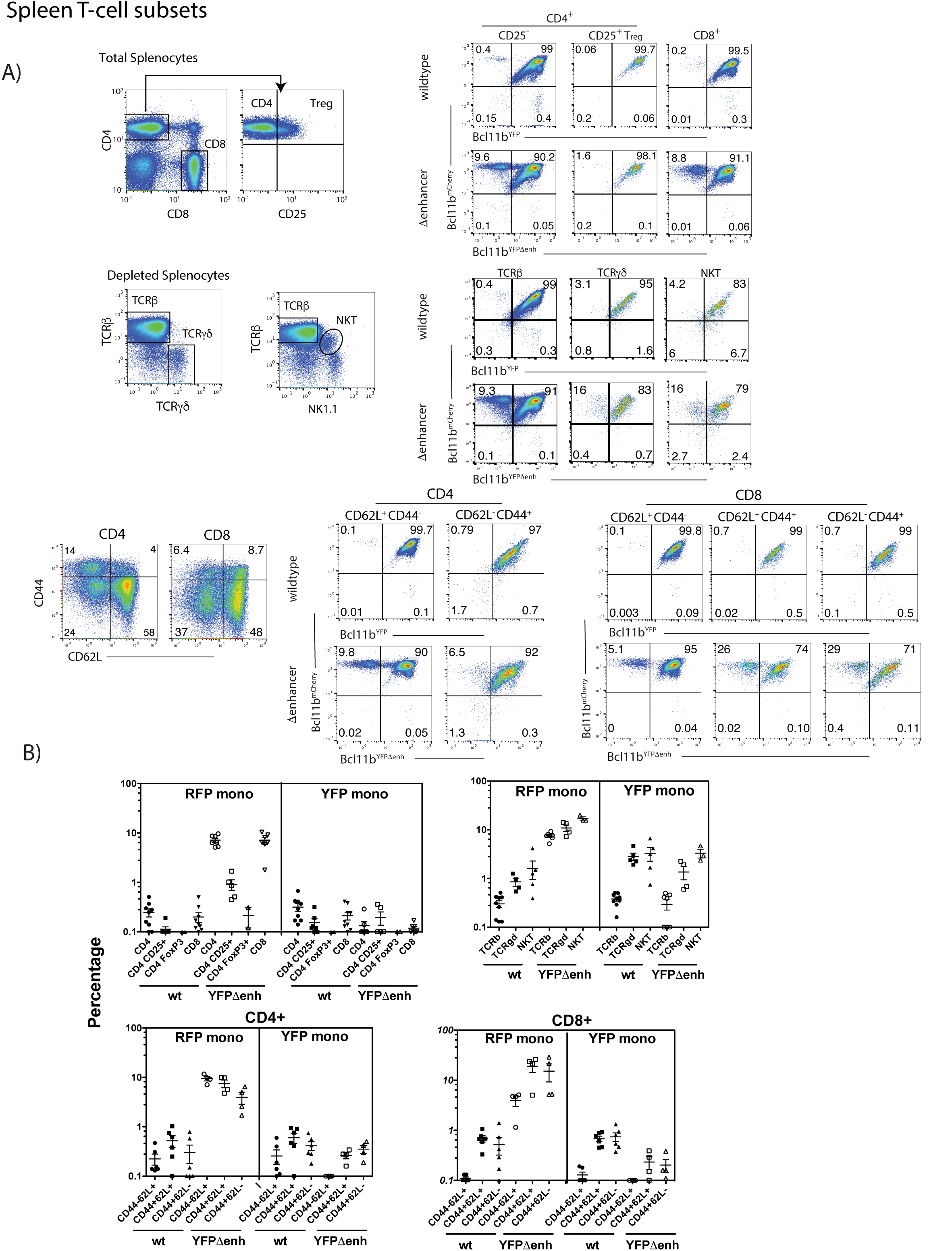
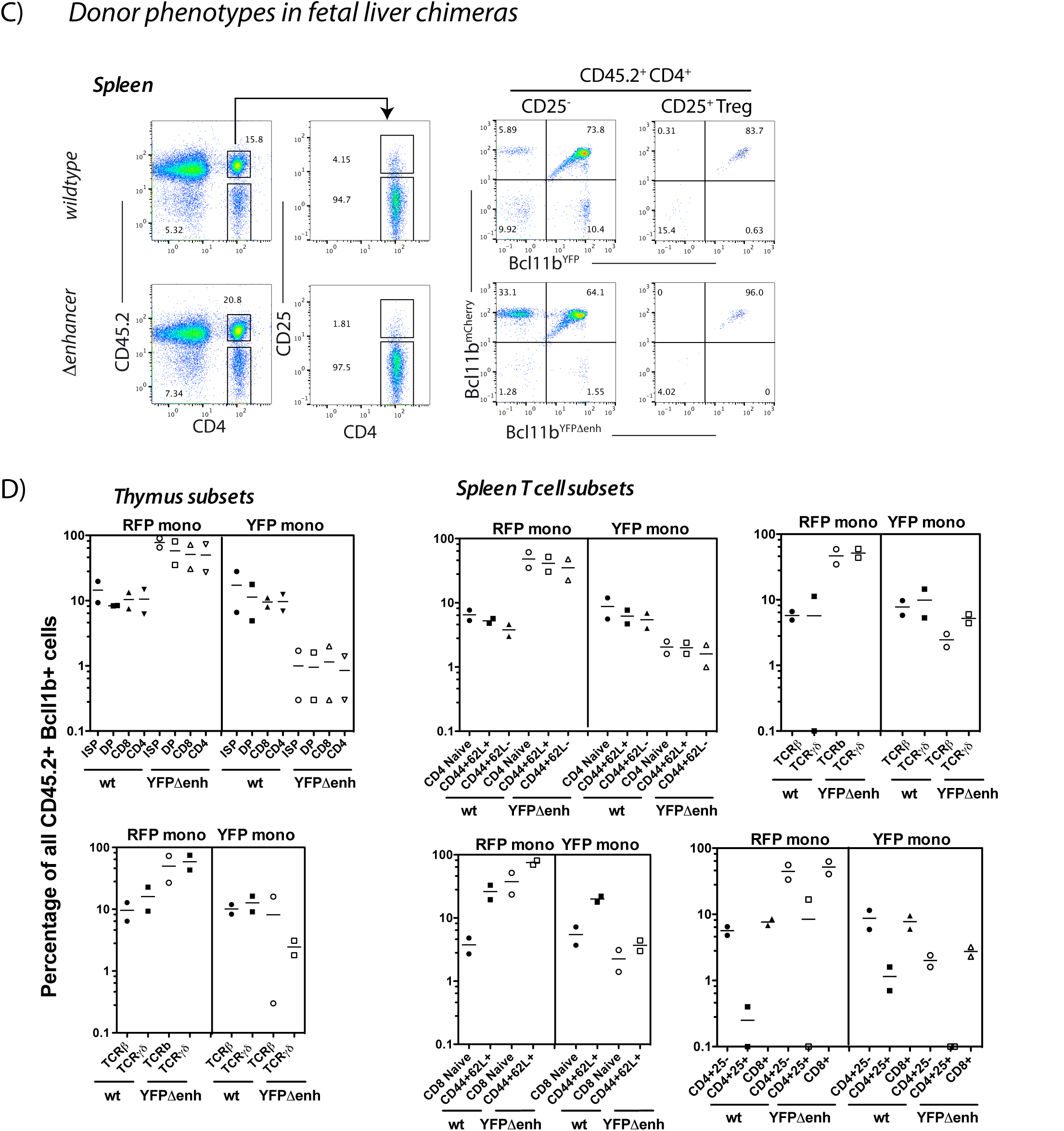
Monoallelic *Bcl11b* expression persists in peripheral splenic T-cell subsets and is cell autonomous. Related to Figure 3. A) Representative flow cytometry plots showing gating strategies for splenic subsets and two-color Bcl11b expression in these populations from Bcl11b^YFP/mCh(neo)^ (wildtype) or Bcl11b^YFPΔEnh/mCh(neo)^ (Enhancer deleted) mice. Some T-cell subsets were enriched by magnetic bead depletion of B cells before staining and analysis as indicated. B) Percentages of cells expressing only mCherry (RFP mono) or YFP (YFP mono) in specific T cell populations from Bcl11b^YFP/mCh(neo^ (wt) or Bcl11b^YFPΔEnh/mCh(neo)^ (YFPΔEnh) mice. Each symbol represents results from an individual mouse (n=2 to 8 mice per group). The data show that patterns of monoallelic expression seen in the thymus (cf. Figure S3) persist in the periphery in CD4, CD8 NKT, and TCRγδ T cells, for both wildtype and YFPΔenh mutant alleles. However, there are subset differences which are most evident in the mCherry wildtype/YFP Δenh genotype. In particular, activated or antigen-experienced (CD44+) CD8 cells show a greater frequency of monoallelic mCherry expression than naïve (CD44-) CD8 cells, whereas CD4+ CD25+ T_reg_ cells exhibit much lower levels of monoallelism than conventional CD4+ and CD8+ cells. These results could be related to the specific requirements for Bcl11b activity in different peripheral T-cell subsets (Avram and Califano, 2014). C)-D) Cell autonomy of Bcl11b expression control in hematopoietic chimeric mice. B6.*Cd45.1* mice were irradiated with 1000 rads and injected retro-orbitally with 10^6^ fetal liver cells from Bcl11b^YFP/mCh(neo)^ (wt) and Bcl11b^YFPΔEnh/mCh(neo)^ (YFPΔEnh) (*Cd45.2+*) mice (F_0_ generation). After 8 weeks chimeric mice were analyzed for expression of the wild type (wt) mCherry and wt or mutant (ΔEnh) YFP alleles. C) Representative flow cytometry plots showing gating strategies for CD45.2+ splenic subsets and two-color Bcl11b expression in these populations from Bcl11b. Other thymic and splenic T-cell populations were gated similarly to those shown Figures S3 and S4. D) Percentages of cells expressing only mCherry (RFP mono) or YFP (YFP mono) in specific T cell populations, demonstrating the persistence of small but similar percentages of monoallelically expressed mCherry and YFP alleles in wt mice and the major increase in monoallelic mCherry positive cells in the presence of the YFPΔEnh mutant alllele. Each symbol represents results from an individual mouse (n=2 mice per group). Results are shown for chimeras from one wildtype/mutant F_0_ donor pair. Similar results were obtained from chimeras from a different pair of wildtype and mutant fetal F_0_ donors.

**Figure S5:**
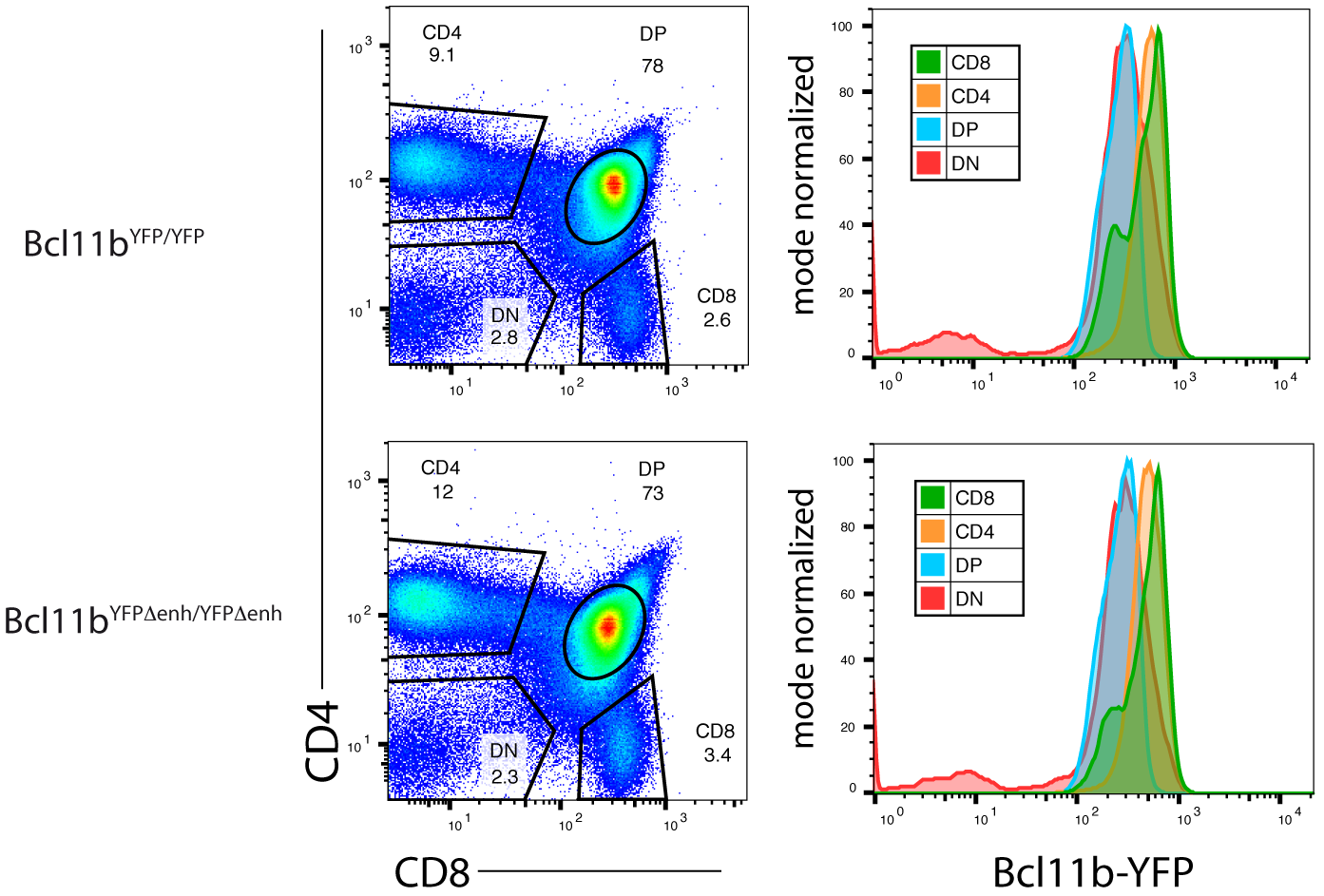
Thymocytes from homozygous mutant enhancer mice Bcl11b^YFPΔEnh/YFPΔEnh^ mice are able to generate T-cell subsets expressing Bcl11b at normal levels relative to wild type enhancer Bcl11b YFP/ YFP mice, Related to Figure 3. Representative FACS plots showing gates used for CD4 and CD8 double negative (DN), double positive (DP) and single positive (CD4 and CD8) populations (left plots) and the relative levels of Bcl11b-YFP in each subset generated from enhancer mutant and wild type mice (right histograms, n=2 for each genotype).

**Figure S6:**
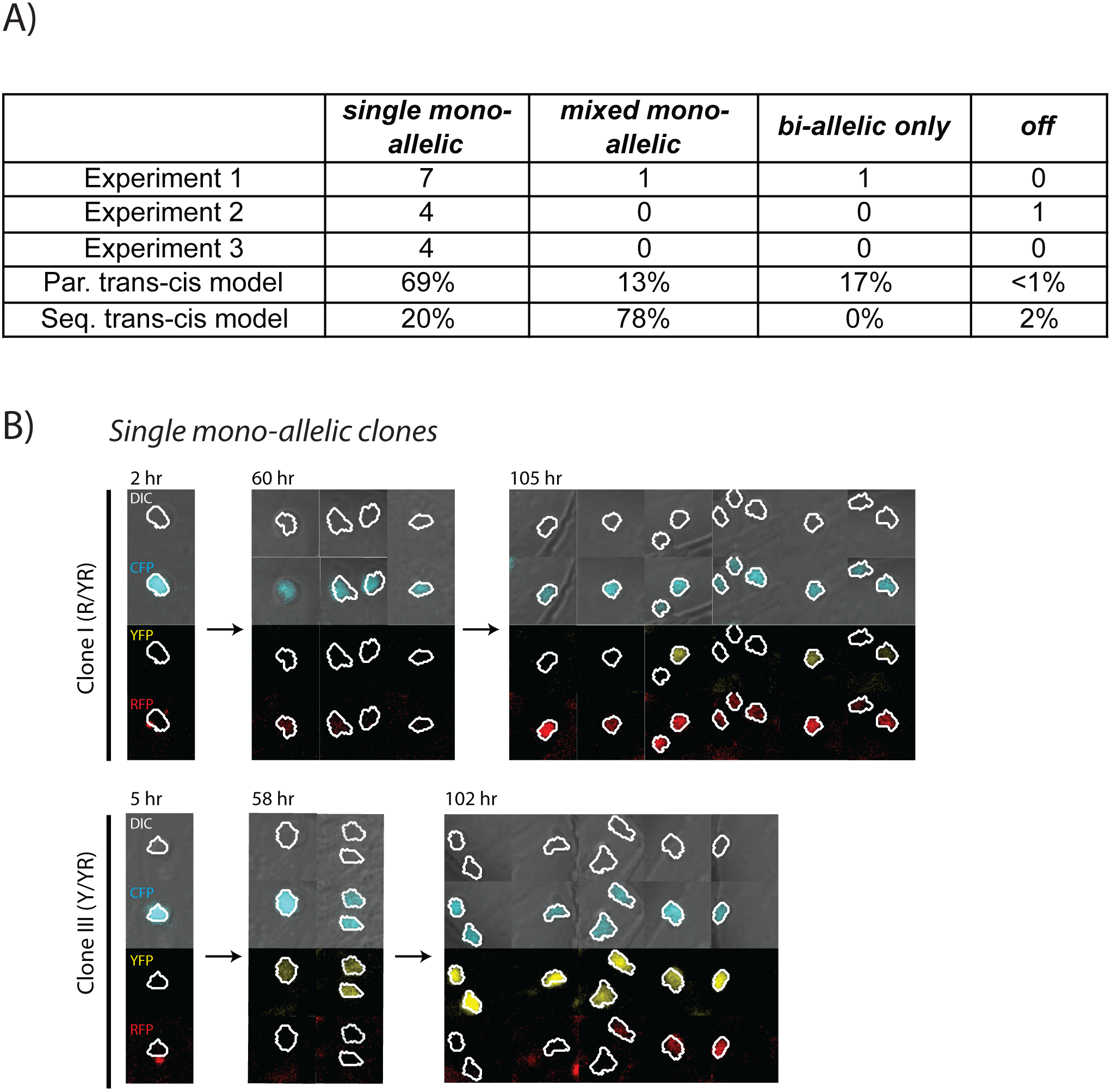
Clones show mono-allelic expression from a single allele during *Bcl11b* activation. A) Table shows observed numbers of clones with indicated allelic activation patterns, showing data from three independent experiments. Simulations of clonal lineages from sequential or parallel *trans-cis* activation models are shown (*N*=30,000 simulations). See Figure 4 and main text for model description and allelic pattern definition. B) Live images show cells from two clonal lineages showing a single mono-allelic pattern of activation, with mono-allelic expression of either the red allele only (Clone I), or the yellow allele only (Clone II). Images shown are from Experiment 3, whereas images and data in Fig. 4 are shown from Experiment 1.

**Figure S7:**
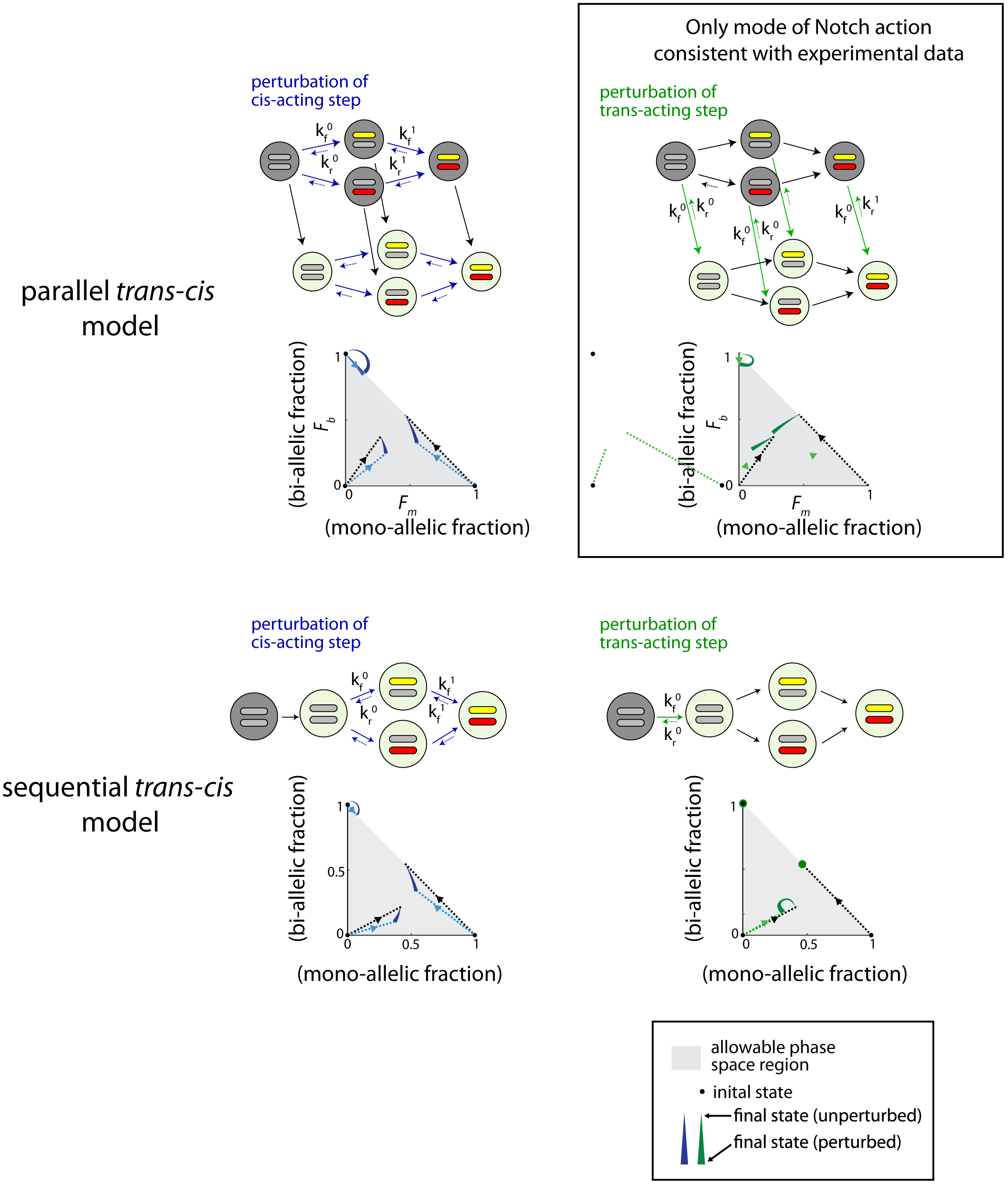
Notch controls a parallel *trans*-acting step for *Bcl11b* activation, Related to Figure 5. The parallel and sequential *trans-cis Bcl11b* activation models (upper and lower panels, respectively). Effects of perturbation of *cis* (blue) or *trans* (green) acting steps in both models are shown with colored arrows. Phase space diagrams show the predicted final fractions of mono-allelic expressing cells (*F_M_*, sum of mono-allelic cells from both alleles), and bi-allelic expressin cells (*F_b_*), for both unperturbed cells (black lines) and cells where the indicated reaction steps are perturbed (colored lines). Definitions for the forward and back rate constants (k_f_^0^, k_f_^1^, k_r_^0^, k_r_^1^) are given in Tables 3-6 of the Mathematical Appendix.

## METHODS

### CONTACT FOR REAGENT AND RESOURCE SHARING

Further information and requests for resources and reagents should be directed to and will be fulfilled by Lead Contact, Ellen Rothenberg

### EXPERIMENTAL MODEL AND SUBJECT DETAILS

#### Animals

F_o_ chimeric mice from Bcl11b^YFP/mCh(neo)^ and Bcl11b^YFPΔEnh/mCh(neo)^ ES-cell blastocyst injections were all made in our lab (described in Method Details). Founder animals were brought to term and crossed in house to generate Bcl11b^YFP(neo)/mCh(neo)^, Bcl11b^YFP/mCh(neo)^, Bcl11b^YFPΔEnh/mCh(neo)^, and Bcl11b^YFPΔEnh/mCh(neo)^ mice. CD45.1 C57BL/6 mice were purchased from Jackson Laboratory. All adult animals were used between 5 and 12 weeks of age. Both male and female mice were used similarly in all studies. Animals were bred and maintained in the Caltech Laboratory Animal Facility, and animal protocols were reviewed and approved by the Institute Animal Care and Use Committee of the California Institute of Technology.

#### Cells

Primary cells isolated from thymus, spleen, bone marrow, and fetal livers were cultured on a OP9-DL1 stromal monolayer system (Schmitt and Zúñiga-Pflücker, 2002) at 37°C in 5% CO_2_ conditions with standard culture medium [80% αMEM (Gibco), 20% Fetal Bovine Serum (Sigma-Aldrich), Pen-Strep-Glutamine (Gibco), 50 μM β-mercaptoethanol (Sigma)] supplemented with appropriate cytokines (described in Method Details).

### METHOD DETAILS

#### Construct Designs

Gene targeting vectors for generating dual allelic *Bcl11b* fluorescent reporter and subsequent enhancer knockout were constructed using a two-step bacterial artificial chromosome (BAC) recombineering method. First, *Bcl11b*-BACs were modified to either insert a fluorescent reporter or disrupt the enhancer sequence with a drug selection marker. An internal ribosome entry site (IRES)-histone 2B-mCherry red fluorescent protein (mCh)-*loxP*-neomycin (*neo*)-*loxP* cassette with homology arms targeting the 3’-untranslated region (UTR) of *Bcl11b* was derived from a similar histone 2B-mCitrine yellow fluorescent protein (YFP) gene targeting vector version published previously (Kueh et al., 2016) and an IRES-H2B-mCherry-*loxP*-neomycin (*neo*)-*loxP* cassette. These two starting plasmids were digested with restriction enzymes NheI and HindIII (New England Biolabs) to exchange the fluorescent protein sequences. Homology arms flanking the 5’ and 3’ ends of the 1.9kb enhancer (Enh) sequence to be replaced (chr12: 108,396,825-108,398,612, mm9 assembly; chr12:107,158,615-107,160,462, in mm10) were attached to a *FRT*-PGK-gb2-hygromycin (*hygro*)-*FRT* drug selection cassette through fusion PCR, and inserted into a cloning vector (pGEM-T-Easy, Promega). Next, restriction enzymes were used to release the homology-flanked fluorescent or drug reporter cassettes, and the resultant linear fragments were introduced into recombineering *E. Coli* strain SW102 containing appropriate BACs for specific targeting. The IRES-mCh-*neo* fragment was linearized with AatII, SalI-HF, ScaI-HF and knocked into a BAC containing the entire *Bcl11b* gene locus (RP24-282D6, from http://bacpac.chori.org). Restriction enyzmes XmnI, PspOMi, and SbfI released the *FRT*-*hygro*-*FRT* cassette used to replace the enhancer sequence in a BAC containing genomic regions downstream of the *Bcl11b* locus (RP23-445J15, from http://bacpac.chori.org). Correctly modified BACs were then selected using kanamycin or hygromycin in combination with chloramphenicol, and verified by PCR and pulse-field gel electrophoresis analysis using the restriction enzyme NotI (New England Biolabs).

A second recombineering reaction retrieved the targeting sequences from reporter modified *Bcl11b*-BACs. The retrieval vector used to fetch the targeting sequence from the modified *Bcl11b*-mCherry-*neo* BAC was made in a previous study (Kueh et al., 2016). For retrieval of the enhancer-disrupted sequence, homology arms for retrieval were first generated using fusion PCR, then cloned into a vector containing a Herpes Simplex Virus-Thymidine Kinase (HSV-TK) cassette using restriction enzymes NotI and SpeI (New England Biolabs). Both retrieval vectors were linearized with PacI and AscI (New England Biolabs), introduced into SW102 containing respective modified Bcl11b-BACs, and retrieved targeting sequences between the homologous ends to generate the desired gene targeting vectors. Clones that underwent correct retrieval reactions were selected using kanamycin or hygromycin in combination with ampicillin, and verified with restriction enzyme digests and sequencing.

The retroviral construct expressing IRES-H2B-mCerulean cyan fluorescent protein (CFP) used for timelapse imaging experiments was generated in a previous study (Kueh et al., 2013). A complete list of vectors used is provided in Key Resources Table.

#### Mouse Generation

A series of genetic modifications were performed to generate different *Bcl11b* reporter mouse strains used for this study (Figure S1). V6.5 mouse embryonic stem (ES) cells with a single modified *Bcl11b* allele expressing the IRES-H2B-mCitrine-*loxp*-*neo*-*loxp* fluorescent reporter were first transfected with Cre recombinase to excise the neomycin cassette. Subclones of this line with a correct deletion of the neomycin cassette were then targeted with the IRES-mCherry-*neo* gene targeting vector to generate dual allelic Bcl11b fluorescent reporter cells, and targeted again with the ΔEnh-*hygro* cassette to delete the enhancer in one allele. After each targeting event, recombinant ES cells grown on feeders were positively selected with antibiotics according to the cassette inserted, and negatively selected with G418. Resistant clones were passaged onto feeder-free conditions and screened using PCR and qPCR for correct targeting. Clones with the desired genotype were karyotyped for normal chromosome numbers before being injected into C57BL/6 blastocyst embryos or subjected to subsequent gene targeting.

F_0_ chimeric mice from *Bcl11b*^YFP/mCh(neo)^ and *Bcl11b*^YFPΔEnh/mCh(neo)^ ES-cell blastocyst injections were generated, and either analyzed at embryonic day 14.5 (E14.5) or brought to term for breeding. *Bcl11b*^YFP/mCh(neo)^ F_0_ chimeric mice were crossed to C57BL/6 mice, and the offspring containing *Bcl11b*-IRES-mCherry-*neo* allele were then bred to homozygosity for this allele. Dual allelic Bcl11b^YFP(neo)/mCh(neo)^ mice with identical *Bcl11b* alleles except for fluorescent protein reporters were generated from breeding *Bcl11b*^mCh(neo)/mCh(neo)^ mice to previously produced *Bcl11b*^YFP(neo)/YFP(neo)^ mice (Kueh et al., 2016), and were used for *in vitro* assay studies of bone marrow derived T-cells. *Bcl11b*^YFPΔEnh/mCh(neo)^ mice were generated in a similar manner by first breeding to C57BL/6 mice to generate enhancer deleted heterozygotes, then crossing mice to *Bcl11b*^mCh(neo)/mCh(neo)^. *Bcl11b*^YFPΔEnh/YFPΔEnh^ mice were generated in parallel by crossing enhancer deleted heterozygotes together. For experiments comparing the effects of the enhancer on *Bcl11b* expression, direct control *Bcl11b*^YFP/mCh(neo)^ mice were generated from breeding *Bcl11b*^YFP/YFP^ and *Bcl11b*^mCh(neo)/mCh(neo)^ animals. However, we have previously reported that the presence or absence of *neo* cassette does not affect the *Bcl11b* reporter locus (Kueh et al., 2016), and do not observe any differences in expression pattern in this study as well (see Figures 1 and 3).

#### Cell Purification

Thymocytes and splenocytes were purified from lymphoid organs removed from 4- to 6- week old normal and enhancer-deleted two-color *Bcl11b* reporter strains, and 2-month post fetal liver precursor transplantation CD45.1 chimeras prior to flow cytometry analysis or fluorescent activated cell sorting (FACS). Harvested lymphoid organs were mechanically dissociated to make single cell suspensions that were re-suspended in Fc blocking solution with 2.4G2 hybridoma supernatant (prepared in the Rothenberg lab). Early stage thymocyte precursors to be analyzed (ETP, DN2a, DN2b, DN3: Figures 1C, 3B, and S3) or sorted (DN2, DN3: Figures 3D and 6A), were first depleted of mature cell lineages using a biotin-streptavidin-magnetic bead removal method. Thymocyte suspensions were labeled with biotinylated lineage marker antibodies (CD8α, TCRβ, TCRγδ, Ter119, Gr-1, CD11c, CD11b, NK1.1), incubated with MACS Streptavidin Microbeads (Miltenyi, Biotec) in HBH buffer (HBSS (Gibco), 0.5% BSA (Sigma-Aldrich), 10 mM HEPES, (Gibco)) pre-filtered through cell separation magnet (BD Biosciences), and passed through a magnetic column (Miltenyi Biotec). Rare T-cell subsets found in the spleen (Figures S4) were enriched using a similar depletion protocol by labeling splenocytes with biotinylated antibodies CD19, CD11b, CD11c, and Gr-1. Later-stage thymocyte precursors analyzed (Figures 1C, 3B, S3, and S5) or sorted (Figures 6A), and whole splenocyte populations analyzed (Figures S4 and S5) were directly stained with conjugated fluorescent cell surface antibodies (see Table S1, Key Resources Table).

Bone Marrow (BM) cells were harvested from dissected femurs and tibiae of 2- to 3- month-old *Bcl11b*^YFP(neo)/mCh(neo)^ mice. Fetal livers (FLs) were removed from F_0_ chimeric fetuses of pregnant surrogate mice at E14.5, individually disrupted mechanically via pipetting into whole organ suspension, and frozen down in freezing media (50% FBS, 40% αMEM, 10% DMSO) for liquid nitrogen storage. Prior to *in vitro* culture use, BM and thawed FL cell suspensions were blocked in 2.4G2 supernatant, tagged with biotinylated antibody lineage markers specific to BM (CD19, CD11b, CD11c, NK1.1, Ter119, CD3ε, Gr-1, B220) or FL (CD19, F4/80, CD11c, NK1.1, Ter119, Gr-1), and depleted of biotin-streptavidin-magnetically labeled mature lineage cells as described above. Eluted lineage depleted (Lin^-^) bone marrow progenitors were either frozen down in freezing media for storage in liquid nitrogen or used directly for *in vitro* cell culture assays of T-cell development, while Lin^-^ fetal liver progenitors were immediately cultured.

#### *In vitro* Differentiation of T-cell Progenitors

DN T cell precursors used for *in vitro* studies were generated by culturing BM and FL stem and progenitor cells on a OP9-DL1 stromal monolayer culture system (Schmitt and Zúñiga-Pflücker, 2002), following previously detailed methods (Kueh et al., 2016) with adapted variations as described below. To promote the DN T cell development, purified or thawed Lin^-^ progenitors were cultured on OP9-DL1-GFP stromal cell monolayers (Schmitt and Zúñiga-Pflücker, 2002) plated on tissue-culture treated plates (Corning) using standard culture medium [80% αMEM (Gibco), 20% Fetal Bovine Serum (Sigma-Aldrich), Pen-Strep-Glutamine (Gibco), 50 μM β-mercaptoethanol (Sigma)], grown at 37°C in 5% CO_2_ conditions, and supplemented with cytokines. All *in vitro* T-cell generation cultures of Bcl11b-YFP/mCh Lin^-^ BM precursors were supplemented with 5 ng/mL Flt-3L (Peprotech) and 5 ng/mL IL-7 (Peprotech), and were sorted after 6 or 7 total days of culture following transduction with a retroviral vector expressing CFP 1 day prior (Figures 2, 4F, 5B, and S2). Lin^-^ fetal liver precursors were cultured with 5 ng/mL Flt-3L and 1 ng/mL IL-7 for the indicated number of days before analysis or sorting.

Sorted thymocytes (Figures 3D and 6A), BM-derived DN2 progenitors (Figure 5B), and FL-DN progenitors (Figure 6B) were seeded manually onto 6000 OP9-DL1-GFP or OP9-Mig feeder cells per well in 96-well plates, cultured in standard medium supplemented with 5 ng/mL Flt-3L and either 5 ng/mL IL-7 (BM) or 1 ng/mL IL-7 (Thymocytes and FL), and harvested for analysis after the indicated number of days.

#### Flow Cytometry and Cell Sorting

Unless otherwise noted, flow cytometry analysis and fluorescent activated cell sorting of all *in vitro* and *ex vivo* lymphocytes were prepared using the procedures outlined. Briefly, cultured cells on tissue culture plates and primary cells from lymphoid organs were prepared as single cell suspensions, incubated in 2.4G2 Fc blocking solution, stained with respective surface cell markers as indicated (see Table S1, Key Resources Table), resuspended in HBH, filtered through a 40-μm nylon mesh, and analyzed using a benchtop MacsQuant VYB flow cytometer (Miltenyi Biotec, Auburn, CA) or sorted with Sony Synergy Sorter (Sony Biotechnology, Inc, San Jose, CA). Both instruments contain capabilities to detect mCherry fluorescence by 561-nm laser excitation. All antibodies used in these experiments are standard, commercially available monoclonal reagents widely established to characterize immune cell populations in the mouse; details are given in Table S1. Acquired flow cytometry data were all analyzed with FlowJo software (Tree Star).

#### Timelapse Imaging

Timelapse imaging of live-cells was used to study *Bcl11b* gene expression dynamics in single cells (Figures 2, 5F, S2, and Movie S1). To prepare for multi-day imaging, PDMS micromesh arrays (250-μm hole diameter, Microsurfaces, AU) containing small microwells that prevent seeded cells from migrating out of a single imaging field of view on 40x objective were adhered to 24 well glass-bottomed plates (Mattek, Ashland, MA). To prevent overcrowding in microwells and enable proper cell tracking, non-GFP expressing OP9-DL1-hCD8 cells (*1*) and sorted CFP+ DN2 progenitors were plated at appropriate densities to achieve ~8 cells/microwell and ~1 cell/microwell, respectively. Cells were cultured in standard medium using Phenol Red-free αMEM (Gibco) and supplemented with 5 ng/mL Flt-3L and 5 ng/mL IL-7.

#### Image segmentation and analysis

Cells were segmented using image processing workflow implemented in MATLAB (Mathworks, Natick, MA), as previously described in detail (Kueh et al., 2013, 2016). Briefly, this workflow involved: 1) Correction for uneven fluorescence illumination, calculated from a fluorescent slide with uniform intensity, followed by background subtraction; 2) Automated cell segmentation, using an Laplacian filter-based edge detection algorithm, followed by exclusion of non cell objects by size and shape selection. Cell segmentations were then subject to manual inspection, and segmented objects that did not correspond to cells were then eliminated. For each data set, automated segmentation parameters were chosen such that the fraction of incorrectly identified cells was <1% of the total number of segmented cells. To calculate fluorescence intensities for segmented cells, we first calculated average intensity levels for an annulus surrounding the segmented cell, and subtracted this background value from image intensities in the cell interior. This additional subtraction was performed to remove auto-fluorescence contributions from OP9-DL1 feeder cells to intensity measurements. Fluorescence intensity measurements were either displayed for clonal cell lineages confined within individual microwells (Figures 2B and S2A), or in a two-dimensional heat map showing the intensity distributions for different indicated time windows for all 218 microwells in a single imaging experiment. To obtain the time evolution of *Bcl11b* population fractions, a least-squares fit of a four-component mixed 2D Gaussian was performed on *Bcl11b* intensity distributions for successive time windows, with each component corresponding to a distinct cellular state (*Bcl11b* YFP^-^mCh^-^, YFP^+^mCh^-^, YFP^-^mCh^+,^ YFP^+^mCh^+^). The relative abundance of each state was then estimated using the area under each best-fit curve (Figure 2D). This approach was used to provide an unbiased estimate of population sizes that did not depend on manually defined thresholds.

#### Model analysis and fitting

Models for *Bcl11b* activation (Models I-III; Figure 4B, Mathematical Appendix) were numerically simulated using an ordinary differential equation solver in MATLAB. The predicted time course from these models were fit to experimental data, using a least-squares procedure with the following free parameters: the *cis*- and *trans*- activation rates (*k_C_* and *k_T_* respectively), and the fraction of cells in each *Bcl11b* non-expressing sub-state, constrained to equal one at *t* = 0 (sequential and parallel *trans-cis* models only). Best fits of the different models were then evaluated by comparing sum-squared errors using the *F*-test, adjusted for different degrees of freedom for each model. Qualitative predictions for perturbing specific reaction steps (Figure S7) were obtained by performing a series of simulations with increasing magnitude of perturbation to the same ending time point. In accordance with experimental observations showing some inactivation of the *Bcl11b* locus upon Notch withdrawal (Figures 5B-C), perturbations involved both a reduction of the forward rate constant, and an increase in the rate of a reverse reaction, together with a graded attenuation in the perturbation after activation of one *Bcl11b* allele (see Figure S7, and Mathematical Appendix for a comprehensive description). Parameters were chosen based on the best fits of the unperturbed time course (Figure 4C), though the direction of the predicted shifts in phase space do not depend on the exact parameters being chosen (Figure 5A).

To generate predictions for allelic state distributions from single clones (Figures 5E-F), we performed Monte-Carlo simulations of clonal single proliferating progenitor lineages, using Markov transition probabilities determined by best-fit rate constants to Models II or III (see Mathematical Appendix). Here, the cell division time was taken to be 20 hrs, corresponding to rates of cell expansion observed in experiments, and measurements of clonal allelic distributions were taken at 100 hrs (i.e. after 5 cell divisions), also matching the time of experimental sampling. Probabilities per cell division for each transition were obtained by converting the continuous-time models to a discrete Markov chain, and these probabilities were taken to be independent between two daughters of the same cell, consistent with the first-order kinetics of these transitions in our models. To test experimental data against each model, we obtained the expected probability of having clones with dual-allelic expression together with monoallelic expression from two alleles (Y+R+D) or from a single allele (Y+D and R+D) clones, for each model, and compared the observed frequencies from clonal lineage data using a chi-squared test.

#### Radiation Chimeras

Fetal liver precursor transplanted CD45.1 chimeras were generated to study the long-term T-cell potential of cells without *Bcl11b* enhancer in mice. Individual fetal liver whole organ suspensions were thawed and split for depletion protocols indicated above or stimulated in standard medium supplemented with 50 ng/mL IL-6 (eBioscience), 50 ng/mL SCF (eBioscience), and 20 ng/mL IL-3 (eBioscience) for 2 days to enrich for hematopoietic stem cell (HSC) progenitors. CD45.1 C57BL/6 mice were subjected to sublethal radiation of 1000 rads from a cesium source. Cells were re-suspended in PBS and 10^6^ cells in a volume of 200 μL were injected retro-orbitally into anesthesized, irradiated mice using 31G, 6 mm insulin syringes (BD). Comprehensive splenocyte analysis was performed on 2-month post transplantation chimeras by sacrificing mice and harvesting spleen and thymus organs following protocols indicated above (Figure S5).

#### Retroviral Transduction on Retronectin-DL1 Coated Plates

Retroviral particles were packaged by transient cotransfection of the Phoenix-Eco packaging cell line with the retroviral construct and the pCL-Eco plasmid (Imgenex) using FuGENE 6 (Promega). Viral supernatants were collected at 2- and 3-days after transfection and immediately frozen at −80°C until use. To infect BM-derived T-cell progenitors, 33 μg/mL retronectin (Clontech) and 2.67 μg/mL of DL1-extracellular domain fused to human IgG1 Fc protein (Varnum-Finney et al., 2000) were added in a volume of 500 μL per well in 24-well tissue culture plates (Costar, Corning) and incubated overnight. Viral supernatants were added next day into coated wells and spun down at 2000 rcf for 2 hours at room temperature. BM-derived T-cell progenitors used for viral transduction were cultured for 5 days according to conditions described above, disaggregated, filtered through a 40-μm nylon mesh, and 10^6^ cells transferred onto each retronectin/DL1-coated virus-bound 24-well supplemented with 5 ng/mL SCF (Peprotech), 5 ng/mL Flt3-L, and 5 ng/mL IL-7.

### QUANTIFICATION AND STATISTICAL ANALYSIS

The sample size for each experiment, and number of independent experiments are stated in the Figures and Figure Legends. In Figure 4C, the best fits of the different models were evaluated by comparing sum-squared errors using the *F*-test (Figure 4D), adjusted for different degrees of freedom for each model. A chi-squared test was applied to compare experimental data against model predictions shown in Figure 4F. Data that had a calculated P-value <0.05 was considered statistical significant, and exact P-values are reported in the figure legends. Bar chart data shown (Figures 6A-B) represent mean and standard deviation.

---

**Table S1:**
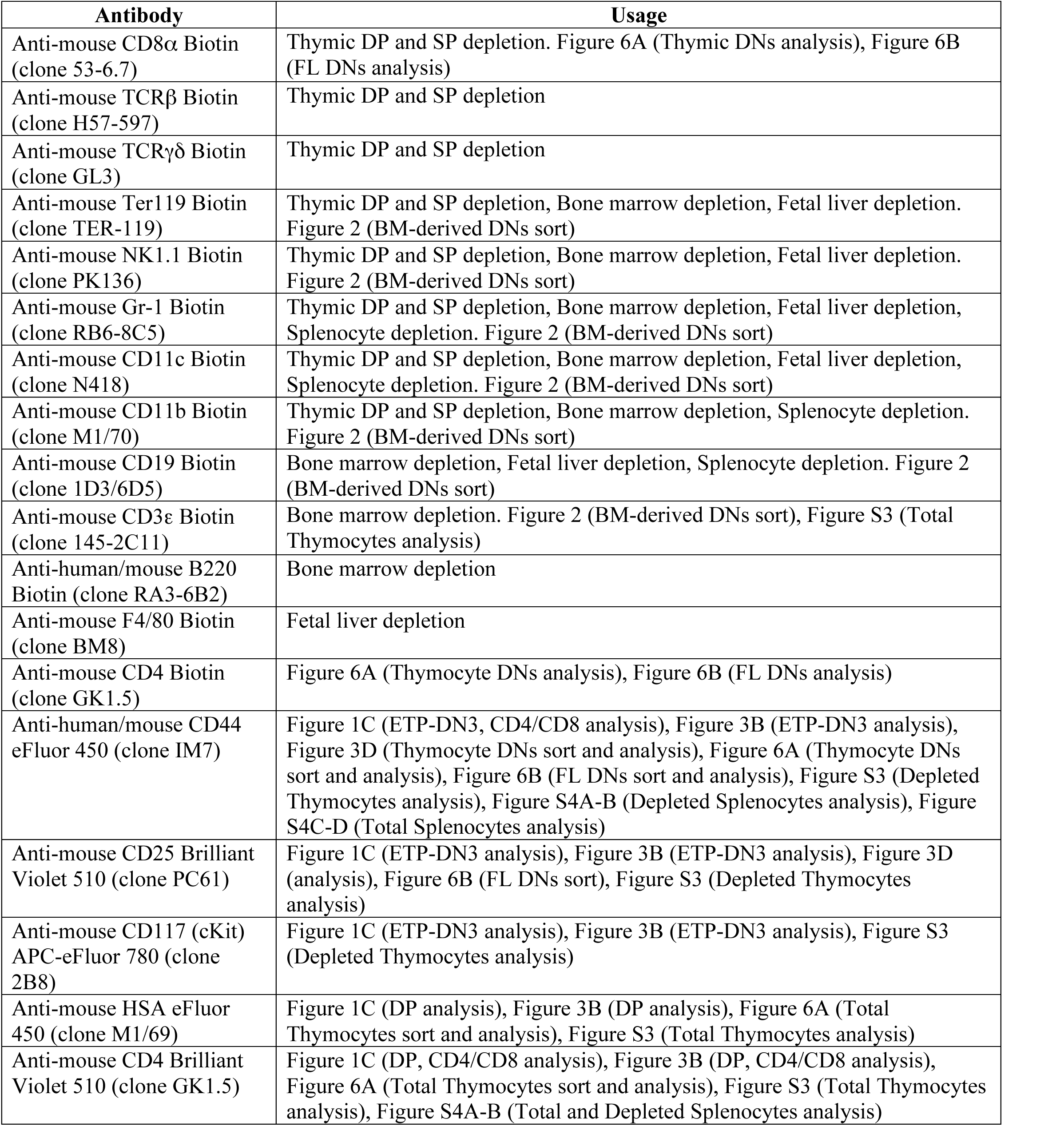
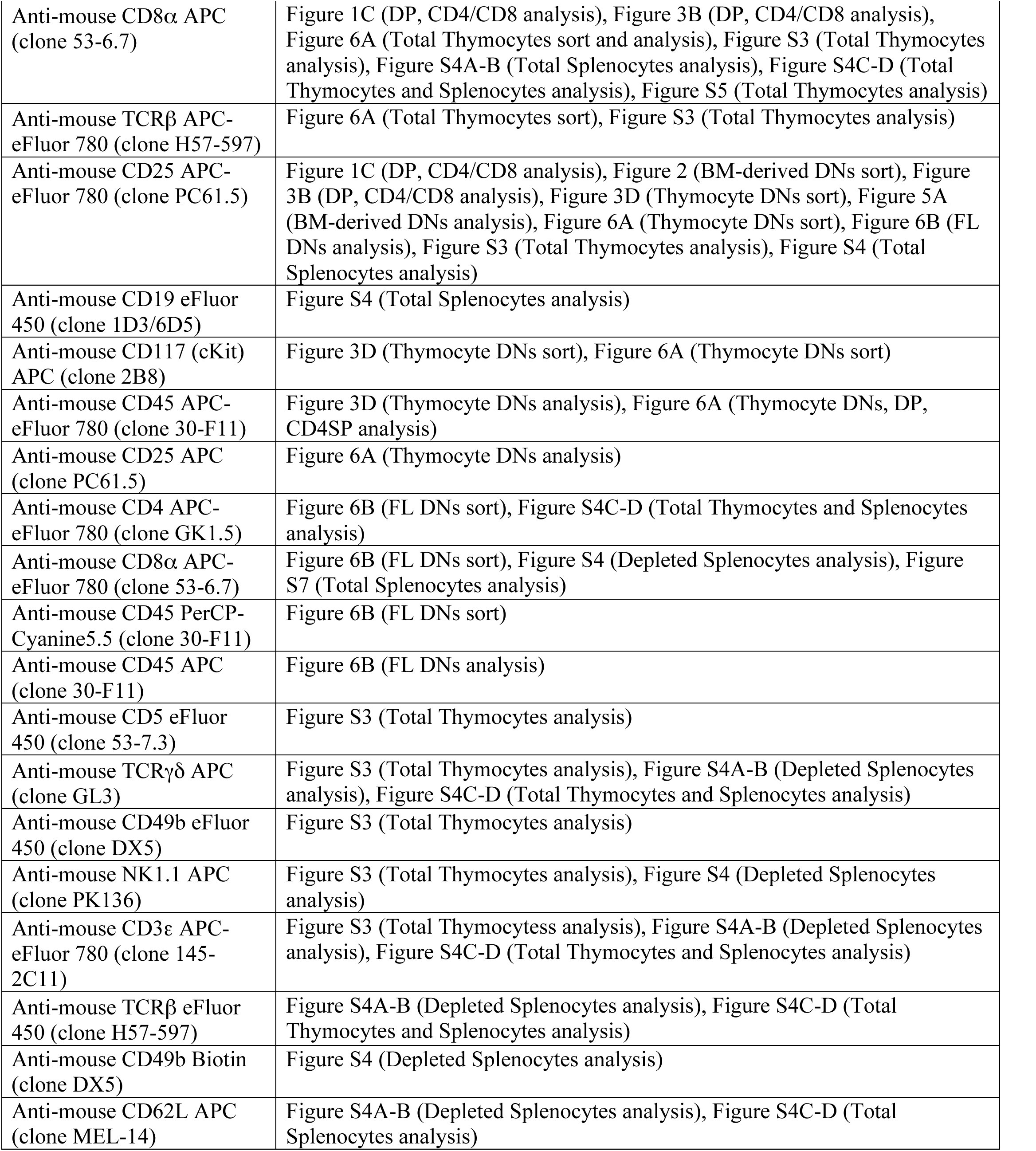
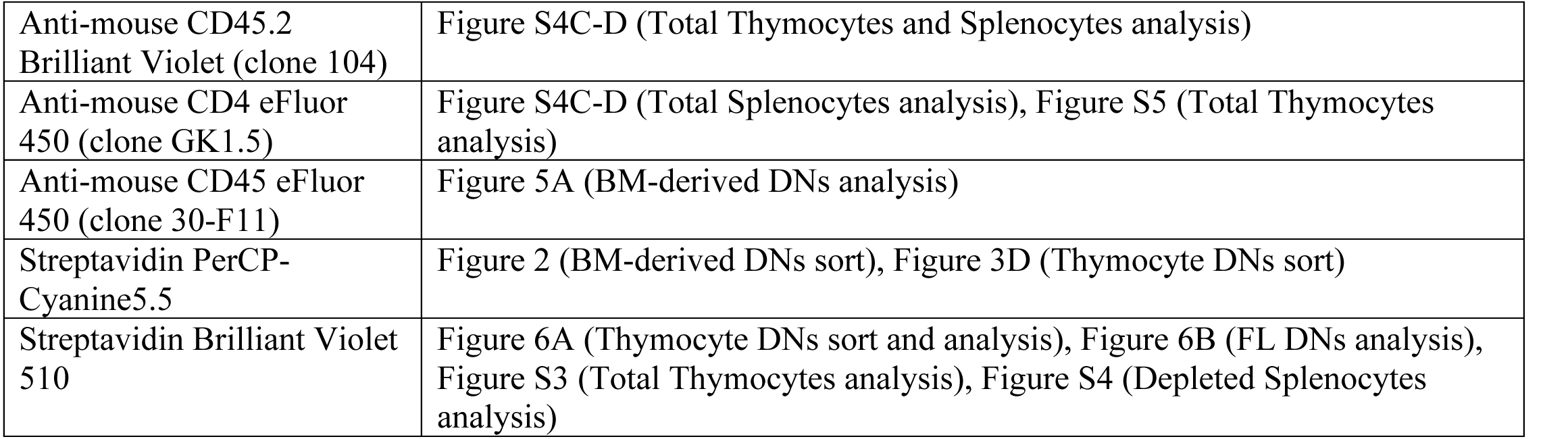
List of Antibodies.

Movie S1: Timelapse movie of clonal DN2 progenitor lineage, Related to Figure 2. Bcl11b-YFP^-^mCh^-^ DN2 progenitors were cultured on OP9-DL1 monolayers with 5 ng/mL IL-7 and Flt3-L within individual PDMS micro-wells, and continuously imaged for 100 hours. Images show superposition of a DIC image (gray) and cellular fluorescent intensities from the Bcl11b-mCherry (red) and Bcl11b-YFP (green) channels, with segmented cell boundaries shown in white. For clarity, images show only the fluorescence intensities within the cell boundaries, excluding auto-fluorescence from well boundaries and OP9-DL1 monolayers. Scatter-plot (bottom-right) updates with each frame to show fluorescent intensities of segmented cells at corresponding time points. Scale bar = 50 microns.

### Mathematical Appendix

We describe a series of dynamical models that aim to clarify the interplay between global (*trans*) and locus-specific (*cis*) mechanisms in the control of *Bcl11b* activation and T-lineage commitment. We first use these models to understand the dynamics of normal *Bc11b* activation in an initial population of DN2 progenitors that are inactive for both *Bcl11b* copies (Figure 4). Next, to distinguish between these different models, we will make predictions about their behavior on a clonal lineage level (Figures 4C-D), and their responses to perturbations of different activation steps (Figures 5C-D and S6, S7), which we will test experimentally. In these models, we do not explicitly model the ETP to DN2 transition, as our experiments all start with cells that have already turned on CD25; however, as we discuss below, our analysis of the sequential and parallel *trans-cis* models suggest that some of the molecular events we consider could occur prior to the ETP-DN2 transition. We note that these models are simplified representations of more complex underlying systems, and a full understanding of the dynamics of the complete system will involve additional processes not accounted for here. However, we use these minimal models to constrain experimental data, evaluate the plausibility of broad classes of mechanisms, and provide a starting point for further investigation.

#### Simple *cis*-activation model

In this model, activation of *Bcl11b* involves a single, slow first-order step that takes place in *cis*, i.e. on the locus of the *Bcl11b* gene itself. This activation step is controlled independently for two copies of *Bcl11b* in a single cell, and with the same rate constant. Under these assumptions, the fraction of non-expressing, monoallelic and biallelic *Bcl11b* expressing cells evolve over time according to the following dynamical equations:

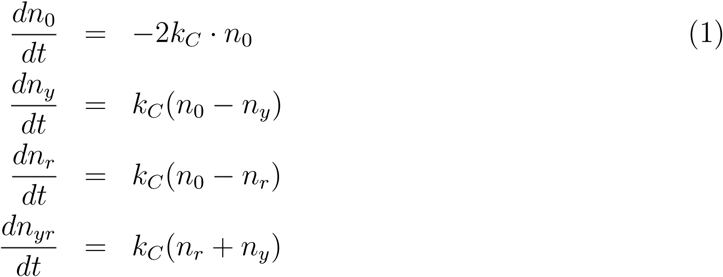

Here *k_C_* is the first-order rate constant of the slow *cis*-acting step on the *Bcl11b* locus. In our experiments, starting DN2 progenitors were sorted to have no *Bcl11b* expression on either copy. Thus, in our model fitting, we take all starting cells to be in a non-expressing state, following this initial condition:

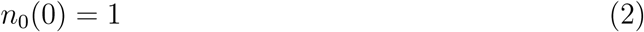

Accordingly, all other variables are set to zero. Following this initial condition, we performed least-squares fitting, varying *k_c_* to provide the best fit to experimental data (Figure 4A). As seen from best least-squares fit, this model is a poor description of the experimentally observed dynamics of *Bcl11b* activation from DN2 (Figures 4A-B): this is because the fraction of biallelic expressing cells increases more slowly compared to that of the monoallelic expressing cells at the earliest time points. To see how this this time lag arises, we can solve for this model analytically, to derive the following solutions:

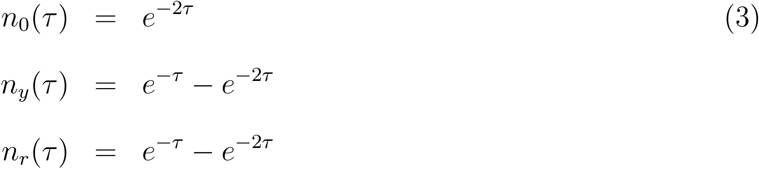

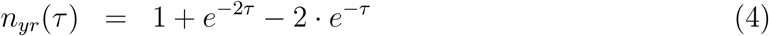

Where *τ* = *k_C_t* is time in non-dimensional units. At early time points, where *τ* ≪ 1, we can expand these solutions using a power series to obtain:

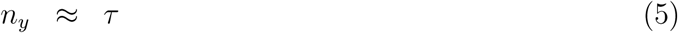

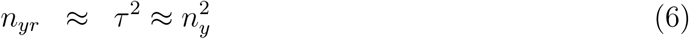

At early time points, the fraction of biallelic expressing cells is approximately the square of fraction of the monoallelic expressing cells, and would therefore increase at a slower rate relative to monoallelic expressing cells.

#### Sequential *trans-cis* activation model

In this model, two rate-limiting steps are required for activation of *Bcl11b*, a *trans*-acting step, which occurs in the nucleus away from the *Bcl11b* locus, and a *cis*-acting step, which occurs independently on each *Bcl11b* locus. The *trans* step precedes, and is necessary for, the *cis* step. Such a model could describe a reaction scheme, where an initial limiting step, occurring away from the *Bcl11b* locus, activates a regulatory factor that facilitates the *cis*-activation step in a permissive fashion. This regulatory factor could be a chromatin-modifying enzyme, a transcription factor, or any other protein that serves to enable locus remodeling. We note that in this model, it is possible that the *trans*-acting step occurs before the DN2 transition (Figure 4A, gray arrows).

There are five states, a *trans*-inactive state *M*_0_, where this *trans* factor is absent, and four states, *N*_0_, *N_y_*, *N_r_*, and *N_ry_*, where the *trans* factor is present, and two copies of *Bcl11b* exist in either active or active states. The time evolution of the fraction of DN2 progenitors in these different states are given by:

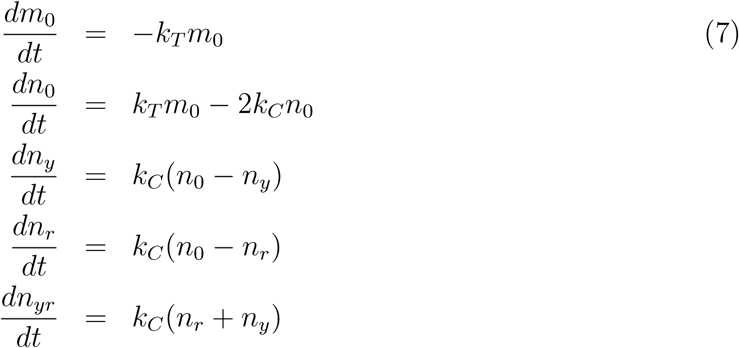

Here, *k_T_* and *k_C_* correspond to the first-order rate constants for the *trans* and *cis*-acting steps respectively. The value of these rate constants were determined using least-squares fitting to experimental data, subject to the constraint that the initial DN2 progenitors that we sorted are all inactive for both copies of *Bcl11b*, and must exist in either *trans*-active or *trans*-inactive states:

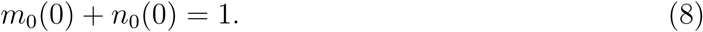

This constraint results in one additional fitting parameter to the model. The best fit trajectory is shown in the main text (Figure 4A), and best-fit parameters are shown in Table 1. Unlike the simple *cis*-activation model, this model can give rise to a rise in the fraction of biallelic expressing fraction concurrent with the monoallelic expressing fraction (Figures 4A-B); thus, from least-squares fitting of experimental data alone, the sequential activation model can plausibly explain experimentally observed population dynamics.

**Table 1:** Best fit parameters of the sequential *trans-cis* activation model to data, with 95% confidence intervals, Related to Figure 4

| Parameter | Units | Best-fit | Lower bound | Upper bound |
| --- | --- | --- | --- | --- |
| $k_c$ | 1/hr | $1.9 \times 10^{-2}$ | $1.5 \times 10^{-2}$ | $2.3 \times 10^{-2}$ |
| $k_t$ | 1/hr | $1.4 \times 10^{-2}$ | $9.1 \times 10^{-3}$ | $1.9 \times 10^{-2}$ |
| $m_0$ | (fraction) | 0.77 | 0.70 | 0.87 |

#### Parallel *trans-cis-*activation model

In this model, *cis*-acting and *trans*-acting steps are also required for activation of *Bcl11b*, similar to the sequential activation model. However, in contrast to the sequential model, *cis* and *trans* steps occur in parallel with each other, such that they occur in either order. In this model, the *trans* step could represent activation of a *trans* factor necessary for transcription of a *cis*-activated locus. For instance, the *trans*-acting step could correspond to the activation of a factor that promotes the polymerase recruitment.

In this model, there are four *trans*-inactive states *M* and four *trans*-active states *N*, each corresponding to different states of locus activation. The time evolution of the fraction of cells in these states are given by:

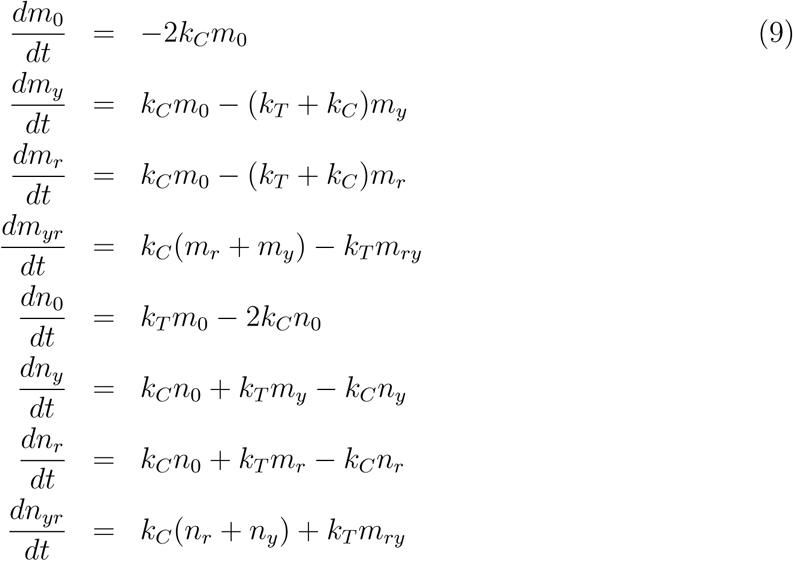

Here, *k_T_* and *k_C_* correspond to the first-order rate constants for the *trans-* and *cis-* acting steps. As experiments start with cells that do not express *Bcl11b*, the following constraint describes the fitting of our models:

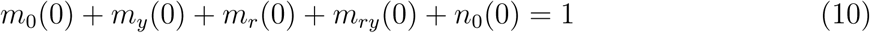

This constraint results in four additional free parameters to the least-squares fit. Upon performing a least-square fit to experimental data, we find that this model also recapitulates the early rise in the fraction of biallelic expressing cells, as observed in the data (Figures 4A-B; see Table 2 for best-fit parameter values). Of note, our model fit suggests that a significant fraction of DN2 progenitors may already exist in a state where one or both *Bcl11b* alleles are already activated in *cis* (Table 2). This feature of our fit will enable us to distinguish between the sequential and parallel activation models using clonal lineage data, as we discuss further below.

**Table 2:** Best fit parameters of the parallel *trans-cis* activation model to data, with 95% confidence intervals, Related to Figure 4

| Parameter | Units | Best-fit | Lower bound | Upper bound |
| --- | --- | --- | --- | --- |
| $k_c$ | 1/hr | $4.2 \times 10^{-3}$ | $8.7 \times 10^{-4}$ | $7.4 \times 10^{-3}$ |
| $k_t$ | 1/hr | $1.4 \times 10^{-2}$ | $1.1 \times 10^{-2}$ | $1.7 \times 10^{-2}$ |
| $m_0$ | (fraction) | 0.21 | 0 | 0.54 |
| $m_r, m_y$ | (fraction) | 0.29 | 0.08 | 0.46 |
| $m_{ry}$ | (fraction) | 0.16 | 0.09 | 0.24 |

#### Comparative analysis of sequential and parallel *trans-cis* activation models

##### Clonal heterogeneity analysis

So far, both sequential and parallel *trans-cis* activation models provide a reasonable fit to the population dynamics of monoallelic and bialellic cell fractions starting from non-expressing progenitors (Figures 4A-B). How can we further distinguish between these two models? So far, we have only considered predictions based on the behavior of whole cell populations; however, analysis of correlations within individual lineage trees can allow discrimination of distinct dynamic mechanisms, as was demonstrated in recent work and in classic experiments [**?**] [**?**] [**?**]. As these two models differ in activation state trajectories taken during *Bcl11b* activation, they would be expected to generate distinct distributions of allelic activation states in single clonal lineages.

To derive *Bcl11b* activation state distributions expected from either sequential or parallel activation models, we first reformulate these models (Eqs. 7 and 9) as discrete time Markov Chains [**?**], where each time step represents a single cell cycle. First, let *N* be the total number of states. Next, define a random state variable *S_t_*, corresponding to the state of the cell at the number *t*. For the sequential model (Eq. 7), the list of states is {*m*_0_*, n*_0_*, n_y_, n_r_, n_yr_*}, with *N* = 5; for the parallel model (Eq. 9), the list of states is {*m*_0_*, m_y_, m_r_, m_yr_, n*_0_*, n_y_, n_r_, n_yr_*}, with *N* = 8. In our descriptions below, we will enumerate all the states as *i* = 1*…N* in such a specified order.

Next, we define *T*, a transition matrix with *N* × *N* elements, where *T_ij_* represents the probability of a cell transitioning from state *j* to state *i* in a single cell cycle. For a given cell cycle time *t_c_*, we can solve the differential equations in Eq. 7 and 9 to obtain corresponding transition probabilities, i.e. *T_ij_*(*t_c_*). In our simulations, we first solve for these transition probabilities, using the best-fit rate constants in Tables 1 and 2. These experiments also used a cell cycle time of *t_c_* = 20 hrs. This was chosen in accordance with the amount of cell expansion observed in imaging experiments, though our conclusions are not expected to depend on the exact value of the cell division time. With this transition probability matrix, we can then simulate state transitions across a lineage of dividing cells, according to the following formula:

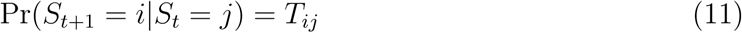

Here *S_t_* represents the state of the cell at the (*t*)th cell cycle. In the Monte-Carlo simulations, each cell gives rise to two cells at each cell division, and each daughter cell chooses its fate randomly and independently from its sibling, based on this formula. This process is repeated iteratively for every descendant from a single ancestor until a designated stopping time (5 cell cycles, or 100 hrs, corresponding to the end of the imaging experiment), whereby a complete lineage tree is generated.

From these clonal lineage simulations, we find the sequential and parallel *trans-cis* activation models yield divergent predictions of heterogeneity in *Bcl11b* allelic activation at the level of single clones. For the sequential activation model, non-expressing ancestors predominantly generate a mixture of progeny with monoallelic expression from both *Bcl11b* copies prior to biallelic *Bcl11b* activation (Figures 4C-D). While some clones only express a single specific *Bcl11b* copy prior to biallelic activation, these clones were rare relative to those with monoallelic expression from each of the two alleles (Figure 4D). This is because all non-expressing progenitors still have both *Bcl11b* copies in a *cis*-inactive state; thus, upon cell division, all daughters of a non-expressing parent retain the same probability of activating either allele.

By contrast, the parallel *trans-cis* activation model gave rise to a large frequency of clones with monoallelic expression from only one specific allele (Figures 4C-D), either red or yellow, varying between, but not within, different clones. This reflects the accumulation of non-expressing progenitors that have a single *Bcl11b* copy present in an open state, but lack the *trans*-acting factors necessary to induce expression from this opened locus (58% total, Table 2). These clones pass through a single specific monoallelic activation intermediate prior to biallelic activation. Additionally, in the parallel activation model, a small percentage of clones transition directly to a biallelic expressing state without first passing through a monoallelic state (Figures 4C-D), a behavior that does not occur for the sequential activation model. This ‘tunneling’ of non-expressing cells to a biallelic expression state reflects the existence of non-expressing cells with both alleles open that still lack the critical *trans*-acting step to enable their expression. For these cells, activation of the *trans*-acting step causes both alleles to turn on simultaneously.

In our experimentally observed distributions of allelic activation states, we found that individual clones predominantly showed monoallelic expression from only one allele (Figure 4E, 7/9 clones observed), but only rarely showed monoallelic expression of both alleles (Figure 4E, 1/9 clones observed). This distribution of single-specific monoallelic clones was significantly different from the fractions predicted for the sequential activation model (*p* < 0.01, *χ*^2^ = 6.8, d.f. = 1), but not significantly different from predictions for the parallel activation model. Furthermore, the experimentally observed distributions also showed evidence for simultaneous activation of both alleles from a non-expressing state (Figure 4E, 1/9 clones), consistent with the occurrence of a parallel *trans*-activation event in a DN2 progenitor with both *Bcl11b* alleles pre-activated in *cis*, which is only allowed in the parallel activation model. Taken together, the experimental clonal lineage data favor the parallel activation model as an explanation for the underlying kinetic processes controlling *Bcl11b* activation, suggesting that the *trans-* acting step necessary for *Bcl11b* activation occurs in parallel with the *cis*-level step on the *Bcl11b* locus.

##### Effects of perturbation of *cis* and *trans* activation steps

To further discriminate between sequential and parallel *trans-cis* activation events, and to gain insights into the molecular mechanisms underlying control of the *trans*-acting step, we analyzed the predicted effects of perturbing different reaction steps for each model. We then tested these predictions by removing the Notch signaling ligand DL1, an essential T-cell developmental signal that controls *Bcl11b* activation probabilities. Here, we show that perturbations of the reaction steps in different models generate distinct shifts in the distribution of monoallelic and biallelic expressing cells, which can be compared to experiments for model discrimination. In this simulation analysis (Figures 5C-D and S7), we perturbed both *cis-*and *trans*-acting steps in the two models in the same way, by reducing its forward rate while introducing a non-zero backward rate. This assumption of reversibility reflects our previous observations that *Bcl11b* can turn back off in a small fraction of cells. We previously noted that although Bcl11b expression maintenance rapidly becomes Notch-independent, there is a small percentage of cells that can lose Bcl11b expression again shortly after activation, if Notch signaling is removed [**?**]. Thus, building on this observation, we reduced the forward rate constant by a fraction *d*, while concomitantly increasing the back rate constant by the same amount *d*. Also, in accordance with experimental observations^1^, the effect of each perturbation on the change in rate constants was further reduced a multiplicative factor *f* (< 1) for transitions to and from a dual-allele expressing state. The perturbed rate constants are labeled in the state transition diagrams in Figure S7, and their definitions, as described here, as listed in Tables 3-6. We note that, while the magnitudes of the experimentally observed shifts depend on these chosen values, the *directions* of these shifts in phase space upon perturbation - corresponding to the increases or decreases in ratio of biallelic to monoallelic expressing cells - are not dependent on the specific values of chosen constants, and thus represents a robust qualitative prediction of the modeling.

**Table 3:**
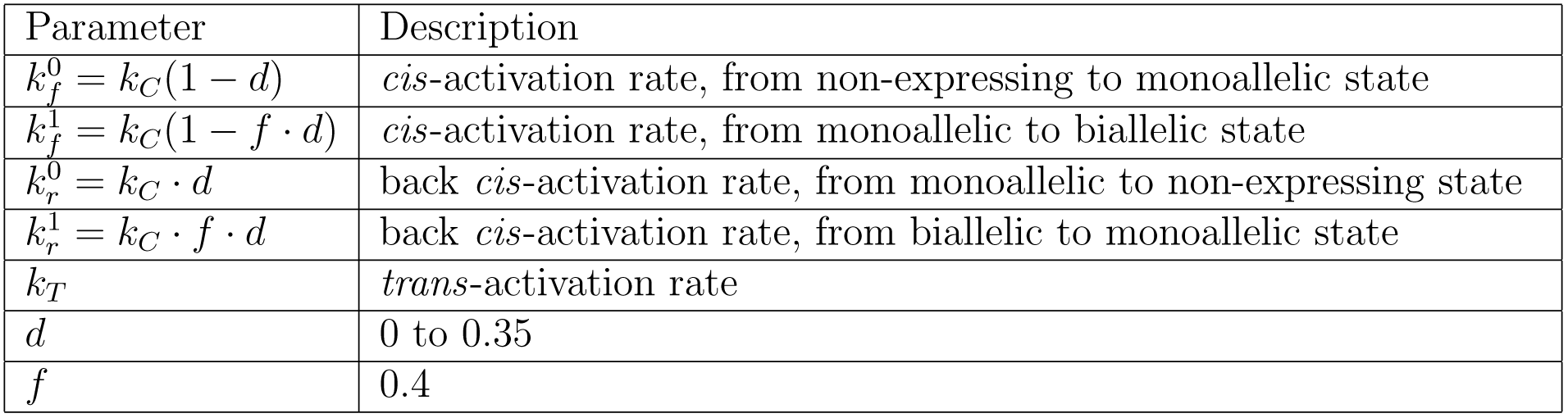
Perturbing the *cis*-acting step in the sequential activation model, Related to Figure 5

**Table 4:** Perturbing the *trans*-acting step in the sequential activation model, Related to Figure 5

| Parameter | Description |
| --- | --- |
| $k_C$ | *cis*-activation rate |
| $k_f^0 = k_T(1 - d)$ | *trans*-activation rate |
| $k_r^0 = k_T \cdot d$ | back *trans*-activation rate |
| $d$ | 0 to 0.35 |

**Table 5:**
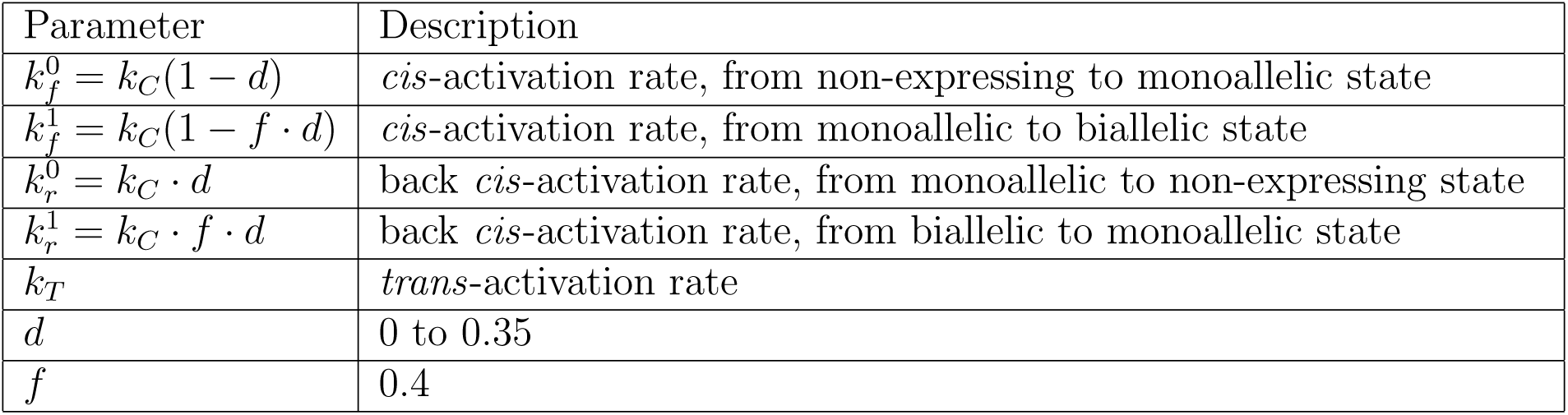
Perturbing the *cis*-acting step in the parallel activation model, Related to Figure 5

**Table 6:** Perturbing the *trans*-acting step in the parallel activation model, Related to Figure 5

| Parameter | Description |
| --- | --- |
| $k_c$ | *cis*-activation rate |
| $k_f^0 = k_T(1 - d)$ | *trans*-activation rate, from non-expressing/monoallelic state |
| $k_f^1 = k_T(1 - f \cdot d)$ | *trans*-activation rate, from biallelic state |
| $k_r^0 = k_T \cdot d$ | back *trans*-activation rate, from non-expressing/monoallelic state |
| $k_r^1 = k_T \cdot f \cdot d$ | back *trans*-activation rate, from biallelic state |
| $d$ | 0 to 0.65 |
| $f$ | 0.2 |

By numerically simulating these models, we found that different perturbations generated distinct shifts in *Bcl11b* monoallelic to biallelic expression states that could then be used to predict the site of Notch action, and distinguish between the two models. Specifically:

- When *cis*-acting steps are perturbed in both the sequential and parallel activation models, non-expressing or monoallelic expressing starting progenitors reach a final state with reduced biallelic expression, and either reduced (sequential activation model) or increased (parallel activation model) monoallelic expression (Figures 5C and S7, blue arrows). However, the ratio of biallelic to monoallelic expressing cells (*F_b_/F_m_*) invariably decreases, such that the line connecting initial to final states in phase space makes a smaller angle with the *x*-axis when perturbation is applied. This qualitative result is not dependent on the exact perturbation strengths specified by *d* and *f* because slowdown of *cis*-acting steps increases the time necessary for cells to achieve full biallelic activation, and thus reduces the amount of biallelic activation at a fixed time.
- When the *trans*-acting step in the sequential activation model is perturbed, progenitors starting without *Bcl11b* expression (Figure S7, sequential model) reach a final state with a reduced fraction of monoallelic and biallelic expressing cells. Progenitors starting with mono or biallelic expression are not affected (Figure S7, sequential model).
- When the *trans*-acting step in the parallel activation model is perturbed, non-expressing and monoallelic progenitors reach a final state with reduced monoallelic and biallelic expression, but also show an increase in the *ratio* of biallelic to monoallelic expressing cells (Figure 5D). As explained above, this increase in *F_b_/F_m_* cannot occur with inhibition of the *cis*-acting step in either model, and cannot occur when starting with monoallelic expressing cells in the sequential activation model. Hence, this shift distinguishes the parallel from the sequential activation model. Furthermore, when starting from a biallelic expressing state, perturbation of the *trans* step causes direct transition to a non-expressing state, without passage through a monoallelic expressing intermediate (Figure S7, parallel model). This biallelic inactivation represents the reversion of the progenitor to a state where cells still maintain two *cis*-active *Bcl11b* alleles *cis*, but have now inactivated a parallel *trans*-acting necessary for expression from a *cis*-opened locus. As this biallelic shutoff would not be predicted to happen upon perturbation of any other step in either model, it provides an additional signature of the parallel activation model.

Taken together, this analysis suggests that the sequential and parallel activation models could potentially be distinguished by analyzing changes in the fractions of non-expressing, monoallelic, and biallelic cells in response to perturbation, if this perturbation involved a disruption of the *trans*-acting step for *Bcl11b* expression.

To test these predictions, we sorted DN2 progenitors with different numbers of active *Bcl11b* alleles, cultured them *in vitro* in either the presence (unperturbed condition) or absence (perturbed condition) of Notch signaling, and then analyzed allelic activation states after four days using flow cytometry. Consistent with parallel activation model, Notch withdrawal reduced the proportion of monoallelic to biallelic expression in cells that started with zero or one active copies of *Bcl11b* (Figures 5A-B). It also caused the direct transition of bialellic expressing cells to a non-expressing state, without passing through a monoallelic intermediate state, as can be seen in the flow cytometry analysis of the effects of Notch removal on progenitors expressing both copies of *Bcl11b* (Figure 5A, green arrow). These experiments reveal a strong correlation at the single-cell level between expression levels of Bcl11b-YFP and Bcl11b-mCherry alleles, in cells that are shutting off their expression, suggesting that the inactivation of *Bcl11b* upon Notch withdrawal occurred in a highly synchronous manner for two alleles. Taken together, our results support the parallel activation model over sequential activation model, and indicate that Notch signaling effectively represents one parallel *trans*-acting step necessary for *Bcl11b* expression.

## Footnotes

1 We attenuated Notch-dependency as described because experiments showed that cells with both *Bcl11b* alleles active show a reduced rate of reversion to an inactive state upon Notch signaling withdrawal. The molecular basis for this attenuation in Notch dependency is currently unclear, but likely involves involve a parallel process occurring in the nuclei of progenitors to stabilize a Notch-driven T-lineage program over time.

